# Experimental estimation of the effects of all amino-acid mutations to HIV Env

**DOI:** 10.1101/067470

**Authors:** Hugh K. Haddox, Adam S. Dingens, Jesse D. Bloom

## Abstract

HIV is notorious for its capacity to evade immunity and anti-viral drugs through rapid sequence evolution. Knowledge of the functional effects of mutations to HIV is critical for understanding this evolution. HIV’s most rapidly evolving protein is its envelope (Env). Here we use deep mutational scanning to experimentally estimate the effects of all amino-acid mutations to Env on viral replication in cell culture. Most mutations are under purifying selection in our experiments, although a few sites experience strong selection for mutations that enhance HIV’s growth in cell culture. We compare our experimental measurements of each site’s preference for each amino acid to the actual frequencies of these amino acids in naturally occurring HIV sequences. Our measured amino-acid preferences correlate with amino-acid frequencies in natural sequences for most sites. However, our measured preferences are less concordant with natural amino-acid frequencies at surface-exposed sites that are subject to pressures absent from our experiments such as antibody selection. We show that some regions of Env have a high inherent tolerance to mutation, whereas other regions (such as epitopes of broadly neutralizing antibodies) have a significantly reduced capacity to tolerate mutations. Overall, our results help disentangle the role of inherent functional constraints and external selection pressures in shaping Env’s evolution.

## Introduction

HIV evolves rapidly: the envelope (Env) proteins of two viral strains within a single infected host diverge as much in a year as the typical human and chimpanzee ortholog has diverged over ~5-million years [1–4]. This rapid evolution is essential to HIV’s biology. Most humans infected with HIV generate antibodies against Env that effectively neutralize viruses from early in the infection [5–7]. However, Env evolves so rapidly that HIV is able to stay ahead of this antibody response, with new viral variants escaping from antibodies that neutralized their predecessors just months before [5–7]. Env’s exceptional evolutionary capacity is therefore central to HIV’s lifestyle.

A protein’s evolutionary capacity depends on its ability to tolerate point mutations. Detailed knowledge of how mutations affect Env is therefore key to understanding its evolution. Many studies have estimated the effects of mutations to Env. One strategy is experimental: numerous studies have used site-directed mutagenesis or alanine scanning to measure how specific mutations affect various aspects of Env’s function [8–17]. However, these experiments have examined only a small fraction of the many possible mutations to Env. Another strategy is computational: under certain assumptions, the fitness effects of mutations can be estimated from their frequencies in global or intra-patient HIV sequences [18–22]. However, these computational strategies are of uncertain accuracy and cannot separate the contributions of inherent functional constraints from those of external selection pressures such as antibodies. Therefore, a more complete and direct delineation of how every mutation affects Env’s function would be of great value.

Recently it has become possible to make massively parallel experimental measurements of the effects of protein mutations using deep mutational scanning [23–25]. These experiments involve creating large libraries of mutants of a gene, subjecting them to bulk functional selections, and quantifying the effect of each mutation by using deep sequencing to assess its frequency pre- and post-selection. Over the last few years, deep mutational scanning has been used to estimate the effects of all single amino-acid mutations to a variety of proteins [26–38]. However, deep mutational scanning has not yet been comprehensively applied to any HIV protein, although it has been used to measure the effects of several thousand single-nucleotide mutations spread across the viral genome [39].

Here we use deep mutational scanning to experimentally estimate how all amino-acid mutations to the ectodomain and transmembrane domain of Env affect viral replication in cell culture. At most sites, our measurements correlate with the frequencies of amino acids in natural HIV sequences. However, there are large deviations at sites where natural evolution is strongly shaped by factors (e.g., antibodies) that are absent from our experiments. Our results show that site-to-site variation in Env’s inherent capacity to tolerate mutations helps explain why some portions of Env evolve so rapidly while other regions are much more conserved. Overall, our work helps elucidate how inherent functional constraints shape Env’s evolution, and demonstrates a powerful experimental approach for comprehensively mapping how mutations affect HIV phenotypes that can be selected for in the lab.

## Results

### Deep mutational scanning of Env

We used the deep mutational scanning approach in Fig 1A estimate the effects of all single amino-acid mutations to Env. We applied this approach to Env from the LAI strain of HIV [40]. LAI is a CXCR4-tropic subtype B virus isolated from a chronically infected individual and then passaged in human T-lymphocytes. We chose this strain because LAI and the closely related HXB2 strain have been widely used to study Env’s structure and function [8–11,41–43], providing extensive biochemical data with which to benchmark our results. LAI’s Env is 861 amino acids in length. We mutagenized amino acids 31-702 (throughout this paper, we use the HXB2 numbering scheme [44]). We excluded the N-terminal signal peptide and the C-terminal cytoplasmic tail, since mutations in these regions can alter Env expression in ways that affect viral infectivity in cell culture [45–47]. The region of Env that we mutagenized spanned 677 residues, meaning that there are 677 × 63 = 42, 651 possible codon mutations, corresponding to 677 × 19 = 12, 863 possible amino-acid mutations.

**Fig 1.**
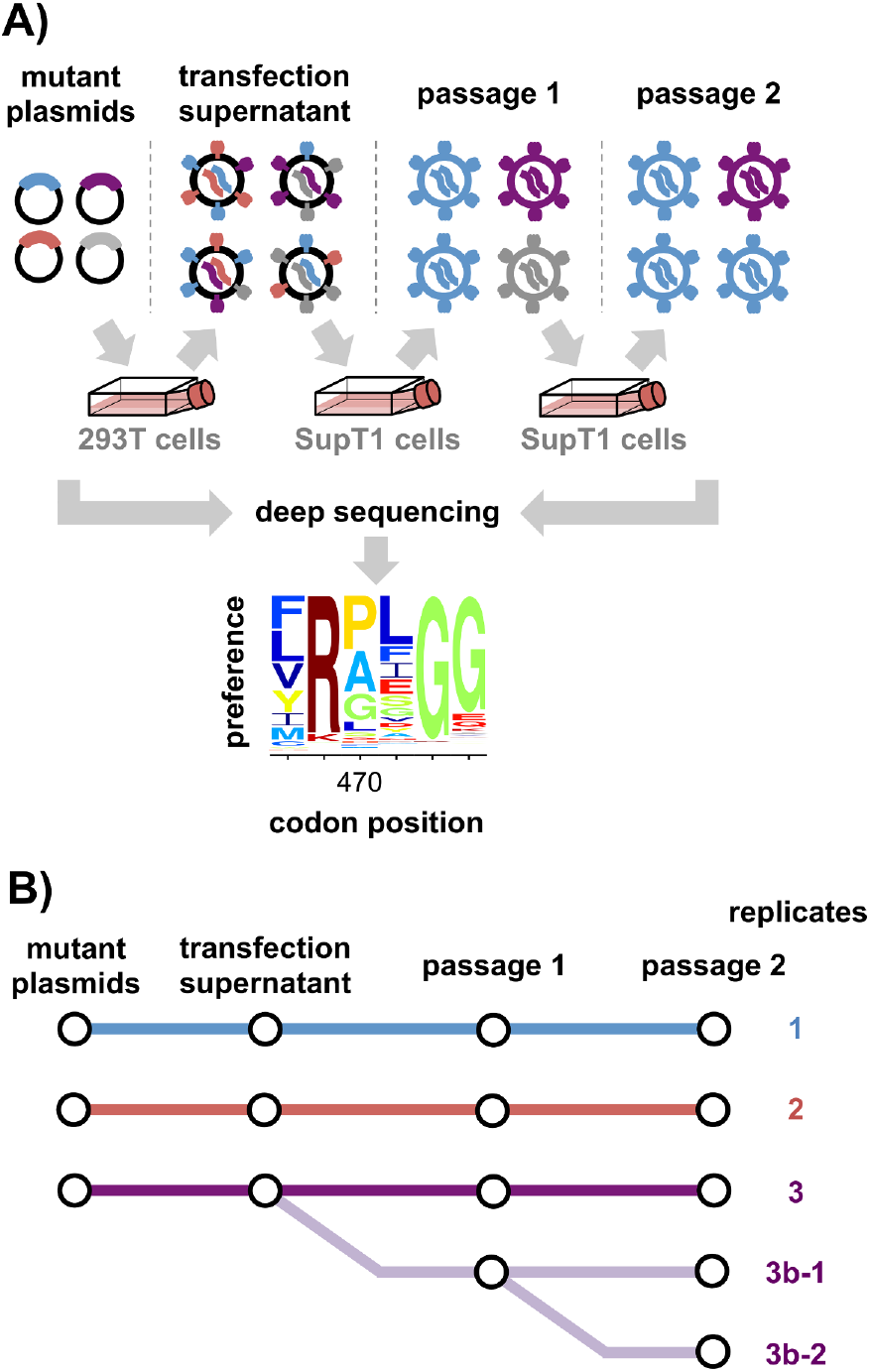
Deep mutational scanning workflow. (**A**) We created libraries of HIV proviral plasmids with random codon mutations in env, and generated mutant viruses by transfecting these plasmid libraries into 293T cells. Since cells receive multiple plasmids, there may not be a link between viral genotype and phenotype at this stage. To establish this link and select for functional variants, we passaged the viruses twice at low multiplicity of infection (MOI) in SupT1 cells. We deep sequenced *env* before and after selection to quantify the enrichment or depletion of each mutation, and used these data to estimate the preference of each site for each amino acid. Each mutant library was paired with a control in which cells were transfected with a wildtype HIV proviral plasmid to generate initially wildtype viruses that were passaged in parallel with the mutant viruses. Deep sequencing of these wildtype controls enabled estimation of the rates of apparent mutations arising from deep sequencing and viral replication. (**B**) We performed the entire experiment in triplicate. Additionally, we passaged the replicate-3 transfection supernatant in duplicate (replicate 3b). We also performed the second passage of replicate 3b in duplicate (replicates 3b-1 and 3b-2).

To create plasmid libraries containing all these mutations, we used a previously described PCR mutagenesis technique [31] that creates multi-nucleotide (e.g, `gca→CAT`) as well as single-nucleotide (e.g, `gca→gAa`) codon mutations. We created three independent plasmid libraries, and carried each library through all subsequent steps independently, meaning that all our measurements were made in true biological triplicate (Fig 1B). We Sanger sequenced 26 clones to estimate the frequency of mutations in the plasmid mutant libraries (S1 Fig). There were an average of 1.4 codon mutations per clone, with the number of mutations per clone roughly following a Poisson distribution. The deep sequencing described in the next section found that at least 79% of the ≈ 10^4^ possible amino-acid mutations were observed at least three times in each of the triplicate libraries, and that 98% of mutations were observed at least three times across all three libraries combined. The plasmid libraries therefore sampled most amino-acid mutations to Env.

We produced virus libraries by transfecting each plasmid library into 293T cells. The viruses in the resulting transfection supernatant lack a genotype-phenotype link, since each cell is transfected by many plasmids. We therefore passaged the transfection supernatants twice in SupT1 cells at an MOI of 0.005 to create a genotype-phenotype link and select for functional variants. Importantly, neither 293T nor SupT1 cells express detectable levels of APOBEC3G [48,49], which can hypermutate HIV genomes [50,51]. This is a crucial point: although HIV encodes a protein that counteracts APOBEC3G, a fraction of viruses will lack a functional version of this protein and so have their genomes hypermutated in APOBEC3G-expressing cells. For each library, we passaged 5 × 10^5^ infectious particles in order to maintain library diversity. We used Illumina deep sequencing to quantify the frequency of each mutation before and after passaging, using molecular barcodes to increase sequencing accuracy as in [37]. We sequenced the plasmids to assess the initial mutation frequencies, and sequenced non-integrated viral DNA [52] from infected SupT1 cells to assess the mutation frequencies in the viruses. Each mutant library was paired with a control in which we generated wildtype virus from unmutated plasmid, enabling us to estimate the rates of sequencing and viral replication errors. Overall, these procedures allowed us to implement the deep mutational scanning workflow in Fig 1.

### Most mutations are under purifying selection, but a few sites experience selection for cell-culture adaptation mutations

Our deep mutational scanning experiments require that selection purge the virus libraries of nonfunctional variants. As an initial gene-wide measure of selection, we analyzed how different types of codon mutations (nonsynonymous, synonymous, and stop-codon mutations) changed in frequency after selection. In these analyses, we corrected for background errors from PCR, sequencing, and viral replication by subtracting the mutation frequencies measured in our wildtype controls from those measured in the mutant libraries (S2 Fig).

Stop-codon mutations are expected to be uniformly deleterious. Indeed, after correcting for background errors, stop codons were purged to <1% of their initial frequency in the twice-passaged viruses for each replicate, indicating strong purifying selection (see the data for “all sites” in Fig 2A). The second viral passage is important for complete selection, as stop codons remain at about ≈16% of their initial frequency in viruses that were only been passaged once (S3 Fig).

**Fig 2.**
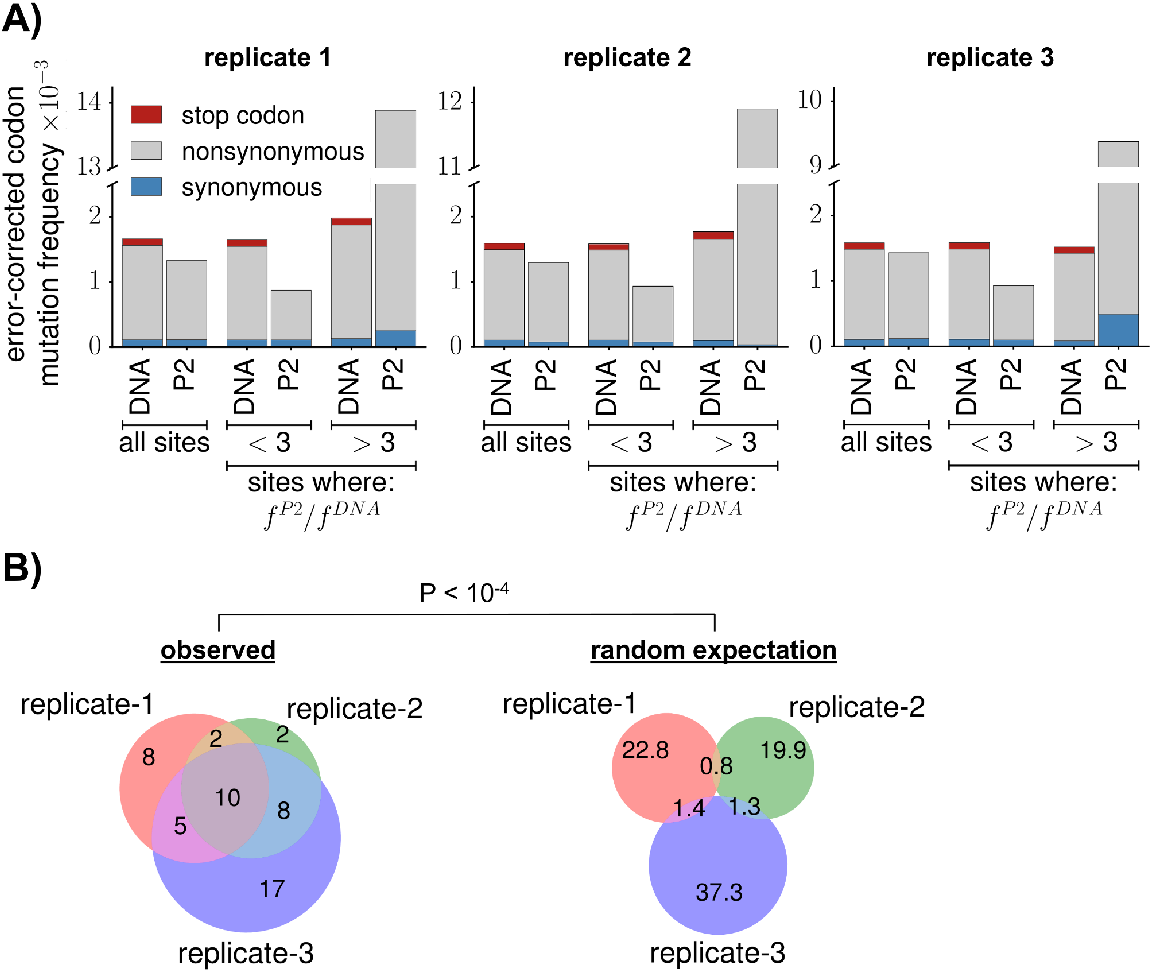
Selection purged mutations in most of env, but favored mutations at a few sites. (**A**) For each replicate, we deep sequenced the initial plasmids (DNA) and the viruses after two rounds of passaging (P2). Bars show the per-codon mutation frequency averaged across sites after subtracting error rates determined from the wildtype controls (S2 Fig). When mutation frequencies are averaged across all sites, selection purged stop codons to <1% of their frequency in the initial DNA. Selection only slightly reduced the average frequency of nonsynonymous mutations; however, this average results from two distinct trends. For ≈4% of sites, the frequency of nonsynonymous mutations in the twice-passaged viruses (*f*^*P*2^) increased >3-fold relative to the frequency in the initial plasmid DNA (*f^DNA^*). For all other sites, the frequency of nonsynonymous mutations decreased substantially after selection. **(B)** The sites at which the error-corrected mutation frequency increased >3-fold are similar between replicates, indicating consistent selection for tissue-culture adaptation at a few positions. The left Venn diagram shows the overlap among replicates in the sites with a >3-fold increase. The right Venn diagram shows the expected overlap if the same number of sites per replicate are randomly drawn from Env’s primary sequence. This difference is statistically significant, with *P* < 10^−4^ when comparing the actual overlap among all three replicates to the random expectation. Another summary view of selection on env is provided by S4 Fig and S5 Fig.

Interpreting the frequencies of nonsynonymous mutations is more nuanced, as different amino-acid mutations have different functional effects. However, a large fraction of amino-acid mutations are deleterious to any protein [53–55]. Therefore, one might expect that the frequency of nonsynonymous mutations would decrease substantially in the twice-passaged mutant viruses. But surprisingly, even after correcting for background errors, the average frequency of nonsynonymous mutations in the passaged viruses is ≈90% of its value in the mutant plasmids (see the data for “all sites” in Fig 2A). However, the average masks two disparate trends. In each library, a few sites exhibit large increases in the frequency of nonsynonymous mutations, whereas this frequency decreases by nearly two-fold for all other sites (see the data for the subgroups of sites in Fig 2A).

An obvious hypothesis is that at a few sites, amino-acid mutations are favored because they are adaptive for viral replication in cell culture. Consistent with this hypothesis, the sites that experienced large increases in mutation frequencies are similar among the three replicates (Fig 2B), suggestive of reproducible selection for mutations at these sites. Moreover, these sites are spatially clustered in Env’s crystal structure in regions where mutations are likely to enhance viral growth in cell culture (Fig 3 and S1 Table). One cluster of mutations disrupts potential glycosylation sites at the trimer apex (Fig 3A), and could be beneficial in cell culture since glycans that shield Env from antibodies in nature [6,56] are often expendable or even deleterious for viral growth in the lab [57–59]. A second cluster overlaps sites where mutations influence Env’s conformational dynamics, which are commonly altered by cell-culture passage [60,61]. A third cluster is at the co-receptor binding interface (Fig 3B), where mutations may enhance viral entry in cell culture. Therefore, while most of Env is under purifying selection against changes to the protein sequence, a few sites are under selection for cell-culture adapting amino-acid mutations.

**Fig 3.**
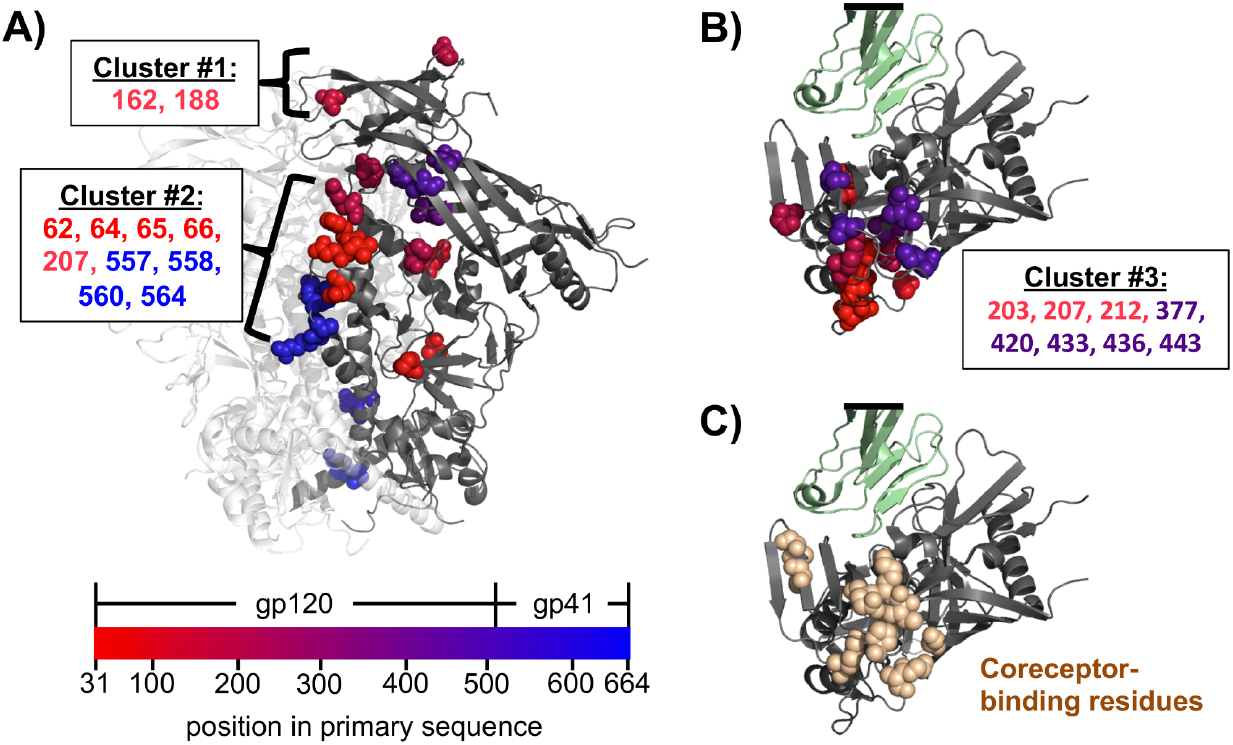
Sites of recurrent cell-culture mutations mapped on Env’s structure. The 25 sites from Fig 2B where the mutation frequency increased >3-fold in at least two replicates after cell-culture passage. **(A)** Trimeric Env (PDB 5FYK [94]) with one monomer in grey and the others in white, oriented so the membrane-proximal region is at the bottom. Sites of cell-culture mutations are shown as spheres, colored red-to-blue according to primary sequence. Most of these sites fall in one of three clusters. Mutations in the first cluster disrupt potential glycosylation sites at Env’s apex. The second cluster includes or is adjacent to sites where mutations are known to affect Env’s conformational dynamics [95,96]. **(B)** The third cluster is near the co-receptor binding surface. This panel shows an apex-down view of monomeric gp120 (grey) in complex with CD4 (green) (PDB 3JWD [42]). Sites of recurrent cell-culture mutations are shown as spheres colored according to primary sequence as in panel A. The black bar indicates cropping of CD4. **(C)** The same view as panel B, but the spheres now show sites known to affect binding to CXCR4 [10] or CCR5 [97]. Note the extensive overlap between the spheres in this panel and panel B.

The average error-corrected frequency of synonymous mutations changes little after selection (an average decrease to 96% of the original frequency; see the data for “all sites” in Fig 2A). This overall trend is consistent with the fact that synonymous mutations usually have smaller functional effects than nonsynonymous mutations. However, synonymous mutations can sometimes have substantial effects [21,62–64], particularly in viruses like HIV that are under strong selection for RNA secondary structure and codon usage [65,66]. To assess selection on synonymous mutations on a more site-specific level, we examined the change in frequency of multi-nucleotide codon mutations across env’s primary sequence (Fig 4). The rationale behind examining only multi-nucleotide codon mutations is that they are not appreciably confounded by errors from PCR, deep sequencing, or *de novo* mutations from viral replication (S2 Fig, S4 Fig). In a region roughly spanning codons 500 to 600, selection strongly purged both synonymous and nonsynonymous multi-nucleotide codon mutations (Fig 4). This region contains *env’s* Rev-response element (RRE) [67], a highly structured region of RNA that is bound by the Rev protein to control the temporal export of unspliced HIV transcripts from the nucleus [68, 69]. The finding of strong selection on the nucleotide as well as the amino-acid sequence of the RRE region of Env therefore agrees with our biological expectations.

**Fig 4.**
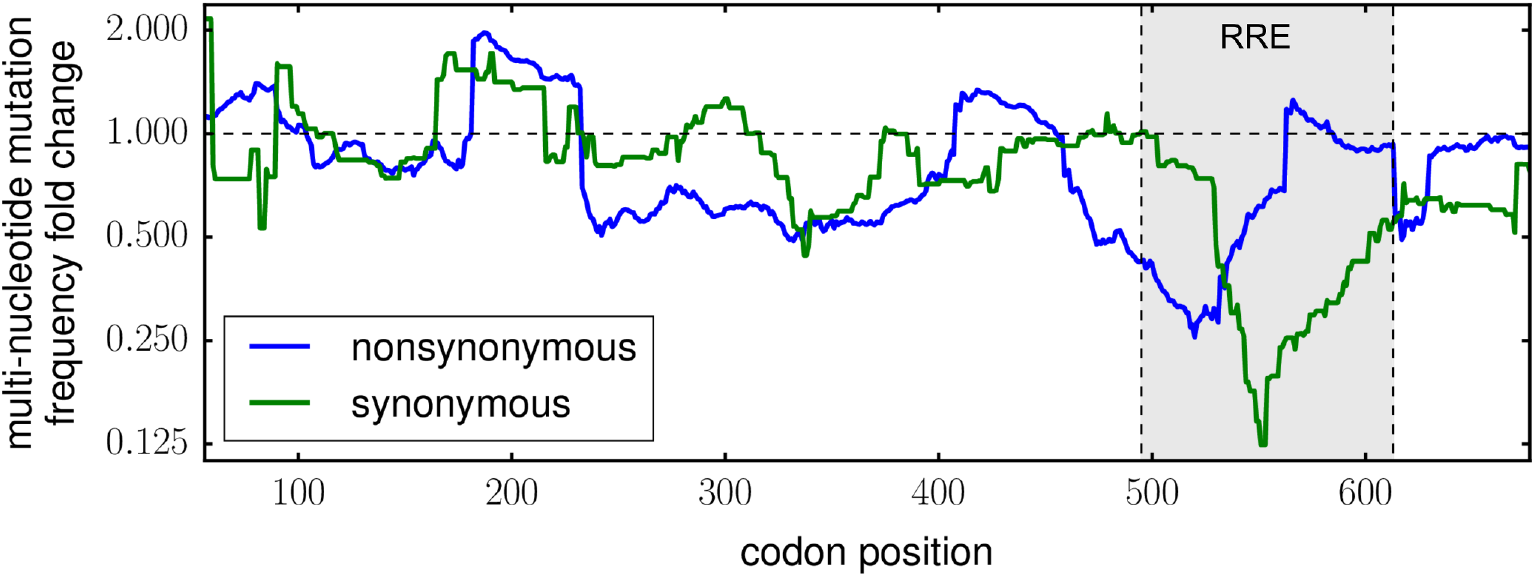
Selection depleted multi-nucleotide codon mutations in the Rev-response element (RRE). This plot shows a 51-codon sliding-window average of the fold change in per-codon multi-nucleotide mutation frequency after two rounds of viral passage, with data plotted for the center point in each window. The strongest depletion of both synonymous and nonsynonymous mutations occurred in the RRE, which is an RNA secondary structure important for viral replication.

### The preference for each amino acid at each site in Env

The previous section examined broad trends in selection averaged across many sites. But our data also enable much more fine-grained estimates of the preference for every amino-acid at every position in Env. We define a site’s preference for an amino acid to be proportional to the enrichment or depletion of that amino acid after selection (correcting for the error rates determined using the wildtype controls), normalizing the preferences for each site so that they sum to one. We denote the preference of site *r* for amino acid *a* as *π_r,a_*, and compute the preferences from the deep-sequencing data as described in [70]. Since we mutagenized 677 residues in Env, there are 677 × 20 = 13, 540 preferences. If selection in our experiments exactly parallels selection in nature and there are no shifts in mutational effects as Env evolves, then these preferences are the expected frequencies of each amino acid at each site in an alignment of Env sequences that have reached evolutionary equilibrium under a mutation process that introduces each amino acid with equal probability [31, 71].

Fig 5 shows Env’s site-specific amino-acid preferences after averaging across replicates and rescaling to account for the stringency of selection in our experiments (details of this re-scaling are in the next section). As is immediately obvious from Fig 5, sites vary dramatically in their tolerance for mutations. Some sites strongly prefer a single amino acid, while other sites can tolerate many amino acids. For instance, site 457, an important receptor-binding residue [8], has a strong preference for aspartic acid. However, this site is adjacent to a variable loop (sites 460-469) where most sites tolerate many amino acids. Another general observation is that when sites tolerate multiple amino acids, they often prefer ones with similar chemical properties. For instance, sites 225 and 226 prefer hydrophobic amino acids, while sites 162 to 164 prefer positively charged amino acids.

**Fig 5.**
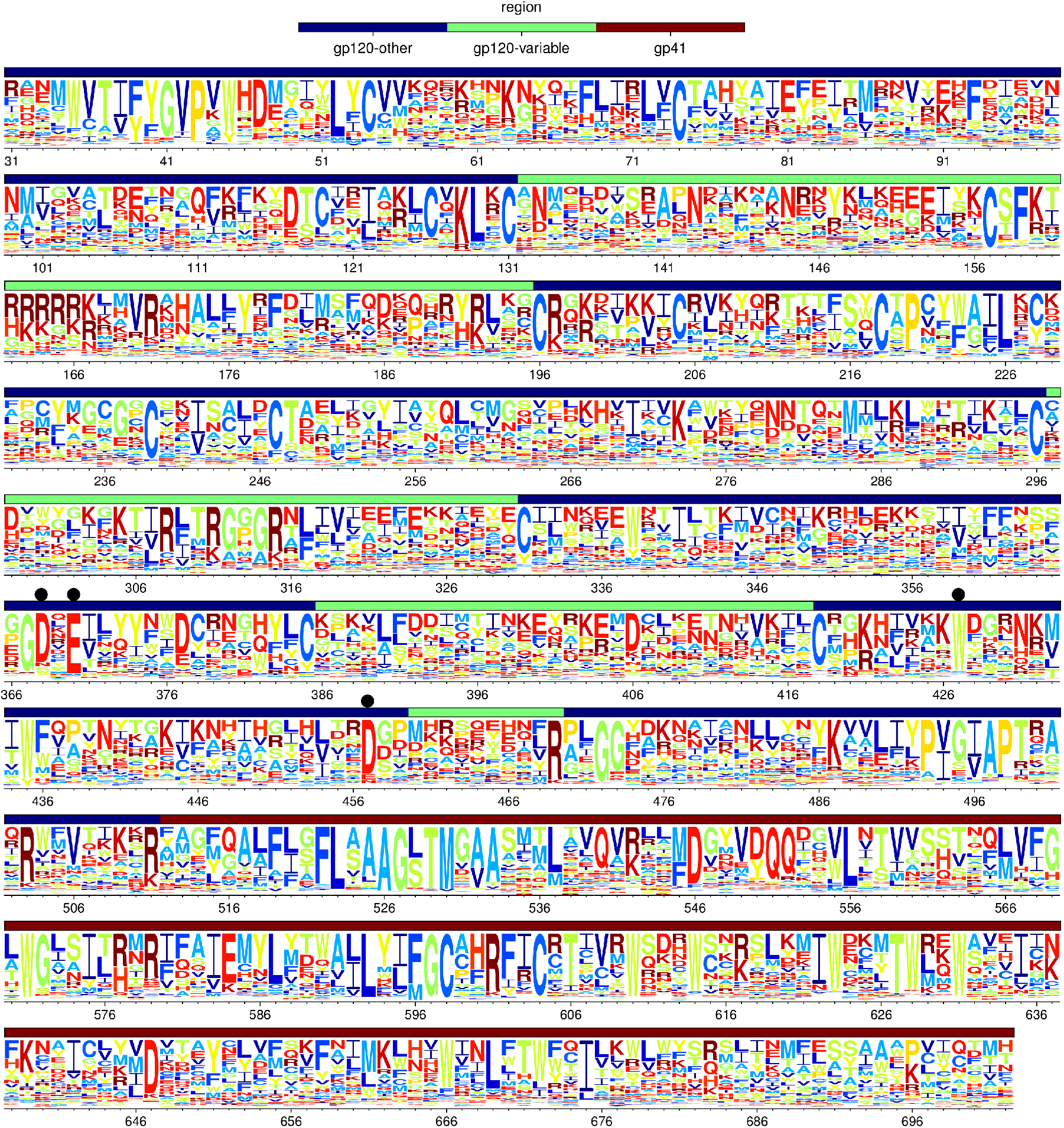
Env’s site-specific amino-acid preferences. The amino-acid preferences averaged across replicates and re-scaled to account for differences in the stringency of selection between our experiments and natural selection. The height of each letter is proportional to the the preference for that amino acid at that site, and letters are colored according to hydrophobicity. The overlay bar indicates the gp120 variable loops, other regions of gp120, and gp41. Black dots indicate sites where mutations are known to disrupt CD4 binding (Table 1). Sites are numbered using the HXB2 scheme [44]. Numerical values of the preferences before and after re-scaling are in S1 File and S2 File, respectively.

To confirm that our experiments captured known constraints on Env’s function, we examined mutations that have been characterized to affect key functions of Env. Table 1 lists mutations known to disrupt an essential disulfide bond, binding to receptor or co-receptor, or protease cleavage. In almost all cases, the deleterious mutation introduces an amino-acid that our experiments report as having a markedly lower preference than the wildtype amino acid. Therefore, our measurements largely concord with existing knowledge about mutations that affect key aspects of Env’s function.

**Table 1.** Our experimental estimates are mostly concordant with existing knowledge about the effects of mutations to functionally or structurally important parts of Env.

| Env function | Site (HXB2 numbering) | Mutation(s) known to disrupt function | Citation | Amino-acid preferences from our experiments |
| --- | --- | --- | --- | --- |
| Disulfide bond | C at 54, 74, 119, 126, 131, 157, 196, 205, 218, 228, 239, 247, 296, 331, 385, 418, 598, 604 | A | [43] | Preference for C at each of these sites is >30-fold higher than for A |
| CD4 binding | D368 | P, R, N, K, E | [8] | Preference for D is >10-fold higher than for these other amino acids |
| CD4 binding | E370 | Q, R | [8] | Preference for E is >100-fold higher than for these other amino acids |
| CD4 binding | W427 | V, S | [8,9] | Preference for W is >100-fold higher than for these other amino acids |
| CD4 binding | D457 | A | [8] | Preference for D is >100-fold higher than for A |
| Co-receptor binding | R298 | A | [10] | Preference for A is actually higher than for R |
| Co-receptor binding | R308 | A | [10] | Preference for R is >100-fold higher than for A |
| Co-receptor binding | R315 | A | [10] | Preference for R is >100-fold higher than for A |
| Co-receptor binding | F317 | A | [10] | Preference for F is >100-fold higher than for A |
| Co-receptor binding | K421 | A | [10] | Preference for K is >100-fold higher than for A |
| Co-receptor binding | Q422 | A | [10] | Preference for A is actually higher than for Q |
| Protease cleavage site | R511 | T | [11] | Preference for R is >100-fold higher than for T |

The preferences listed in the last column are the average from all replicates, re-scaled by the stringency parameter in Table 2.

A crucial aspect of any high-throughput experiment is assessing the reproducibility of independent replicates. Fig 5 shows the *average* of the preferences measured in each replicate. Fig 6A shows the correlations among the 13,540 site-specific amino-acid preferences estimated from each of the three replicates. The correlations are modest, indicating substantial replicate-to-replicate noise. In principle, this noise could arise from differences in the initial plasmid mutant libraries, bottlenecks during the generation of viruses by transfection, bottlenecks during viral passage, or bottlenecks during the sequencing of proviral DNA from infected cells. Analysis of technical replicates of the first or second round of viral passaging indicates that most of the noise arises from bottlenecks during the viral passaging or sequencing steps. Specifically, measurements from replicate 3 are no more correlated to those from replicates 3b-1 or 3b-2 (which are repeated passages of the same transfection supernatant, Fig 1B) than they are to those from totally independent replicates (compare Fig 6 and S6 Fig). However, replicates 3b-1 and 3b-2 (which shared the first of the two viral passages, Fig 1) do yield more correlated measurements than independent replicates (S6 Fig). The existence of bottlenecks during viral passage is also suggested by the data in S4 Fig and S5 Fig. Therefore, the experimental reproducibility could probably be increased by passaging more infectious viruses at each step.

**Fig 6.**
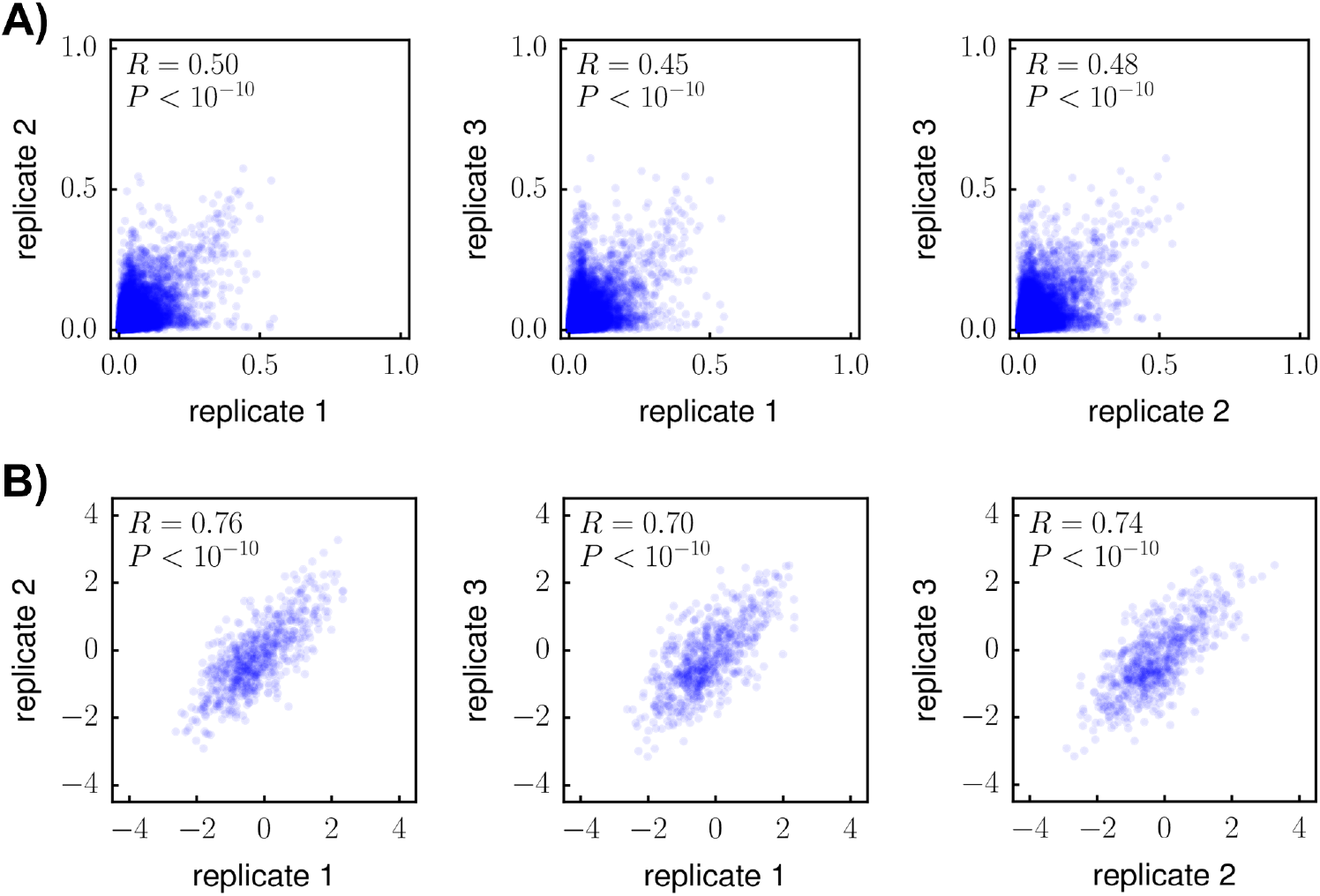
The amino-acid preferences are modestly correlated among experimental replicates, but the preferred amino acids have highly similar hydrophobicities. **(A)** Correlations between the site-specific amino-acid preferences from each replicate. **(B)** Correlations between the preference-weighted hydrophobicities. For each site r, the preference-weighted hydrophobicity is *∑_a_ π_r,a_* × *X_a_* where *π_r,a_* is the preference of *r* for amino acid *a*, and *X_a_* is the Kyte-Doolittle hydropathy [98] of a. The fact that the hydrophobicities are more correlated than the amino-acid preferences means that when different amino acids are preferred at a site in different experimental replicates, the chemical properties of the preferred amino acids are similar. Each plot shows the Pearson correlation coefficient and associated P-value. Similar data for replicates 3b-1 and 3b-2 are in S6 Fig.

If bottlenecks cause each replicate to sample slightly different mutations, then perhaps in each replicate there will be selection for similar amino acids even if the exact mutations differ. To test this hypothesis, we quantified the extent that each site preferred hydrophobic or hydrophilic amino acids. We did this by computing a site-specific hydrophobicity score from the amino-acid preferences. Fig 6B shows that these preference-weighted hydrophobicities are more correlated between replicates than the preferences themselves. Therefore, even though there is replicate-to-replicate noise in the exact amino acids preferred at a site, the preferred amino acids have similar chemical properties among replicates.

### The amino-acid preferences correlate with amino-acid frequencies in HIV sequence alignments at most sites, but deviate at positions subject to selection pressures absent from our experiments

In the previous section, we showed that our experimentally measured amino-acid preferences captured the constraints on Env’s biological function for sites with known mutational effects (Table 1). If this is true across the entire protein, then our measurements should correlate with the frequencies of amino acids in natural HIV sequences. Table 2 shows that there is a modest correlation (Pearson’s R ranging from 0.29 to 0.36) between the preferences from each experimental replicate and the frequencies in an alignment of HIV-1 group-M sequences (a phylogenetic tree of these sequences is in Fig 7A). Since each replicate suffers from noise due to partial bottlenecking of the viral diversity, we hypothesized that averaging the preferences across replicates should make them more accurate. Indeed, averaging the replicates increased the correlation to *R* = 0.4 (Table 2).

**Fig 7.**
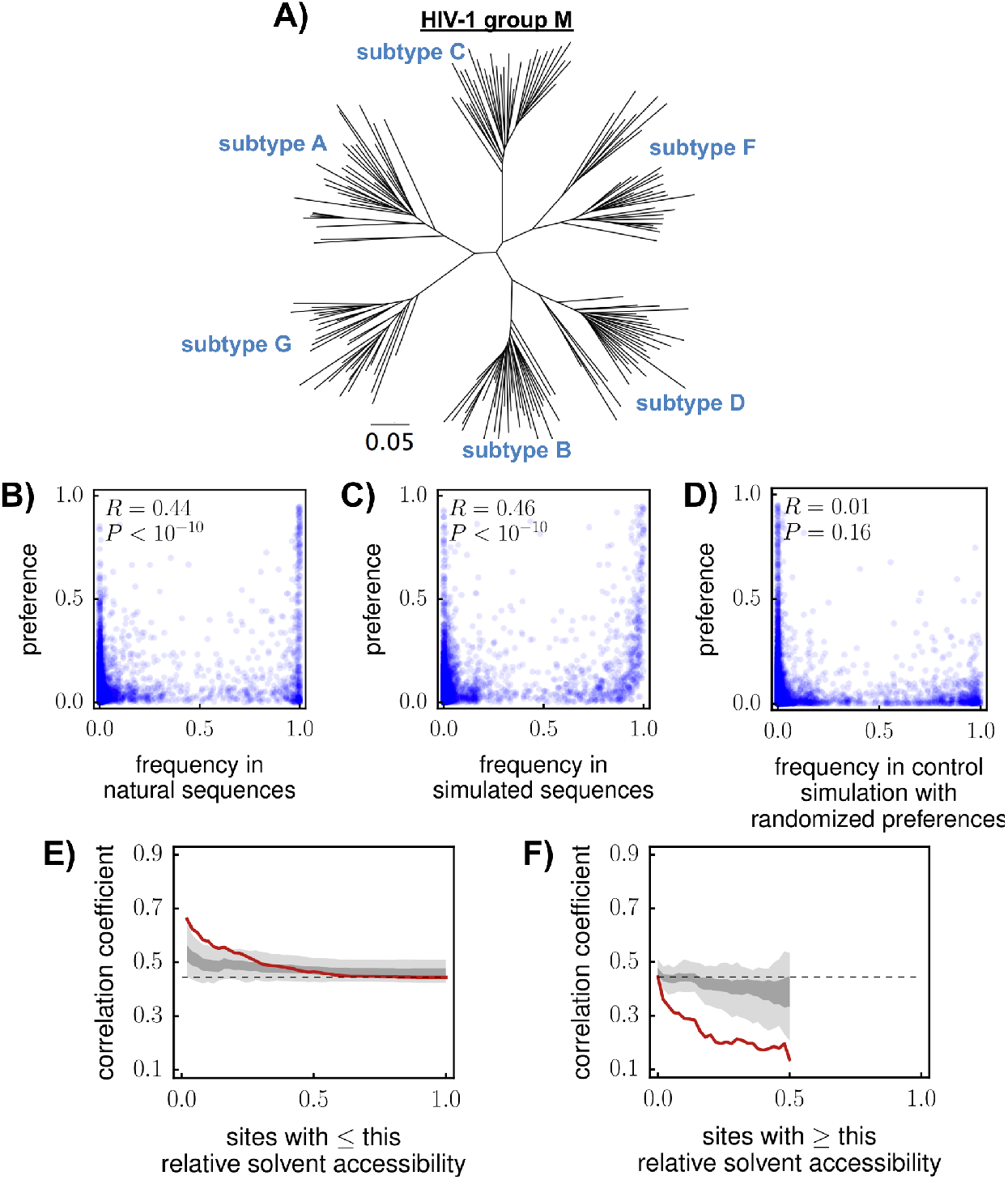
Correlations between amino-acid preferences and frequencies in natural HIV sequences. **(A)** Phylogenetic tree of the HIV-1 group-M sequences in the alignment. **(B)** Correlation between alignment frequencies and preferences. The preferences are the replicate averages re-scaled by the stringency parameter in Table 2. **(C)** The correlation if evolution is simulated along the phylogenetic tree assuming that the preferences correctly describe the actual selection. **(D)** There is no correlation in a control simulation in which preferences are randomized among sites. **(E), (F)** Correlation between preferences and alignment frequencies as a function of relative solvent accessibility (RSA). Red lines show the actual correlation. Dark and light gray show the range of correlations in the middle 80% and 100% of 100 simulations. For both plots, data are shown until the subset of sites that meets the RSA cutoff becomes less than 10% of all sites in Env; this is why neither x-axis extends all the way from 0 to 1. Correlation coefficients are Pearson’s R.

**Table 2.** Correlation of amino-acid preferences with amino-acid frequencies in nature.

| replicate | correlation | | stringency parameter ($\beta$) |
| --- | --- | --- | --- |
|  | preferences | rescaled preferences |  |
| 1 | 0.32 | 0.33 | 1.7 |
| 2 | 0.31 | 0.32 | 1.6 |
| 3 | 0.29 | 0.29 | 1.4 |
| 3b-1 | 0.36 | 0.37 | 1.5 |
| 3b-2 | 0.35 | 0.36 | 1.5 |
| average | 0.40 | 0.44 | 2.1 |

Pearson correlation between experimentally measured amino-acid preferences and frequencies of amino acids in an alignment of HIV-1 group-M sequences. Correlations are shown for both raw preferences and preferences re-scaled by the stringency parameter that maximizes the correlation. The correlation is highest when the preferences are averaged across replicates and re-scaled by a stringency parameter $> 1$ .

The concordance between deep mutational scanning measurements and natural sequence variation is improved by accounting for differences in the stringency of selection in the experiments compared to natural selection [71,72]. Specifically, if the measured preference is *π_r,a_* and the stringency parameter is *β*, then the re-scaled preference is 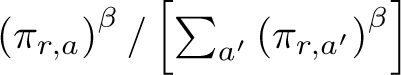. A stringency parameter of *β* > 1 means that natural evolution favors the same amino acids as the experiments, but with greater stringency. Table 2 shows that for all replicates, the stringency parameter that maximizes the correlation is > 1. Therefore, natural selection prefers the same amino acids as our experiments, but with greater stringency.

After averaging across replicates and re-scaling by the optimal stringency parameter, the Pearson correlation is 0.44 between our experimentally measured preferences and amino-acid frequencies in the alignment of naturally occurring HIV sequences (Fig 7B). Is this a good correlation? At first glance, a correlation of 0.44 seems unimpressive. But we do not expect a perfect correlation even if the experiments perfectly concord with selection on Env in nature. The reason is that natural HIV sequences are drawn from a phylogeny (Fig 7A), not an ideal ensemble of all possible Env sequences. The frequencies of amino acids in this phylogeny reflect evolutionary history as well as natural selection. For instance, if several amino acids are equally preferred at a site, one is likely to be more frequent in the alignment due to historical contingency. Additionally, natural evolution is influenced by the genetic code and mutation biases: a mutation from the tryptophan codon `TGG` to the valine codon `GTT` is extremely unlikely even if valine is more preferred than tryptophan. Therefore, the correlation will be imperfect even if the preferences completely concord with natural selection –the question is how the actual correlation compares to what is expected given the phylogenetic history and mutation biases.

To determine the expected correlation, we simulated evolution along the phylogenetic tree in Fig 7A under the assumption that the experimentally measured preferences exactly match natural selection. Specifically, we used `pyvolve` [73] to simulate evolution using the experimentally informed site-specific codon substitution models described in [72], which define mutation-fixation probabilities in terms of the amino-acid preferences. In addition to the preferences and the stringency parameter β = 2.1 from Table 2, the substitution models in [72] require specification of parameters reflecting biases in the mutation process. We estimated nucleotide mutation bias parameters of *φ_A_* = 0.55, *φ*_C_ = 0.15, *φ_G_* = 0.11, and *φ_T_* = 0.18 from the frequencies at the third-nucleotide codon position in sequences in the group-M alignment for sites where the most common amino acid had 4-fold codon degeneracy. We used the transition-transversion ratio of *κ* = 4.4 estimated in [74].

The correlation between the preferences and amino-acid frequencies in a representative simulated alignment is shown in Fig 7C. As this plot illustrates, the expected correlation is only about 0.46 if the experimentally measured preferences exactly describe natural selection on Env under our model. As a control, we also simulated evolution using substitution models in which the preferences have been randomized among sites (Fig 7D); as should be the case, there is no correlation in these control simulations. So the actual correlation is nearly as high as expected if natural selection concords with the preferences measured in our experiment.

We next investigated if there are parts of Env for which there is an especially low correlation between our experimentally measured preferences and natural amino-acid frequencies. For instance, antibodies exert selection on the surface of Env in nature [6,7,75,76]. We therefore examined the actual and simulated correlations between the preferences and frequencies as a function of solvent accessibility (Fig 7E,F). For all sites (right side of Fig 7E, left side of Fig 7F), the actual correlation is only slightly lower than the range of correlations in 100 simulations. For more buried sites, both the simulated and actual correlations increase (Fig 7E), presumably because sites in the core of Env tend to have stronger preferences for specific amino acids. But as sites become more surface-exposed, the actual correlation drops below the value expected from the simulations (Fig 7F). Therefore, our experiments provide a relatively worse description of natural selection on Env’s surface than its core – probably because the evolution of the protein’s core is shaped mostly by inherent functional constraints that are effectively captured by our experiments, whereas the surface is subject to selection pressures (e.g., antibodies) that are not modeled in our experiments.

Comparing disulfide-bonded cysteines and glycosylation sites vividly illustrates this dichotomy between inherent functional constraints and external selection pressures. Env has 10 highly conserved disulfide bonds, most of which are essential for the protein’s inherent function [43]. Env also has numerous N-linked glycosylation sites, many of which are also highly conserved in nature, where they help shield the protein from antibodies [6,56]. Fig 8 shows that our experimentally measured preferences are highly correlated with natural amino-acid frequencies at the sites of the disulfides, but not at the glycosylation sites. This result can easily be rationalized: the disulfides are inherently necessary for Env’s function, whereas the glycosylation sites are important largely because of the external selection imposed by antibodies. Our experiments therefore accurately reflect the natural constraints on the former but not the latter.

**Fig 8.**
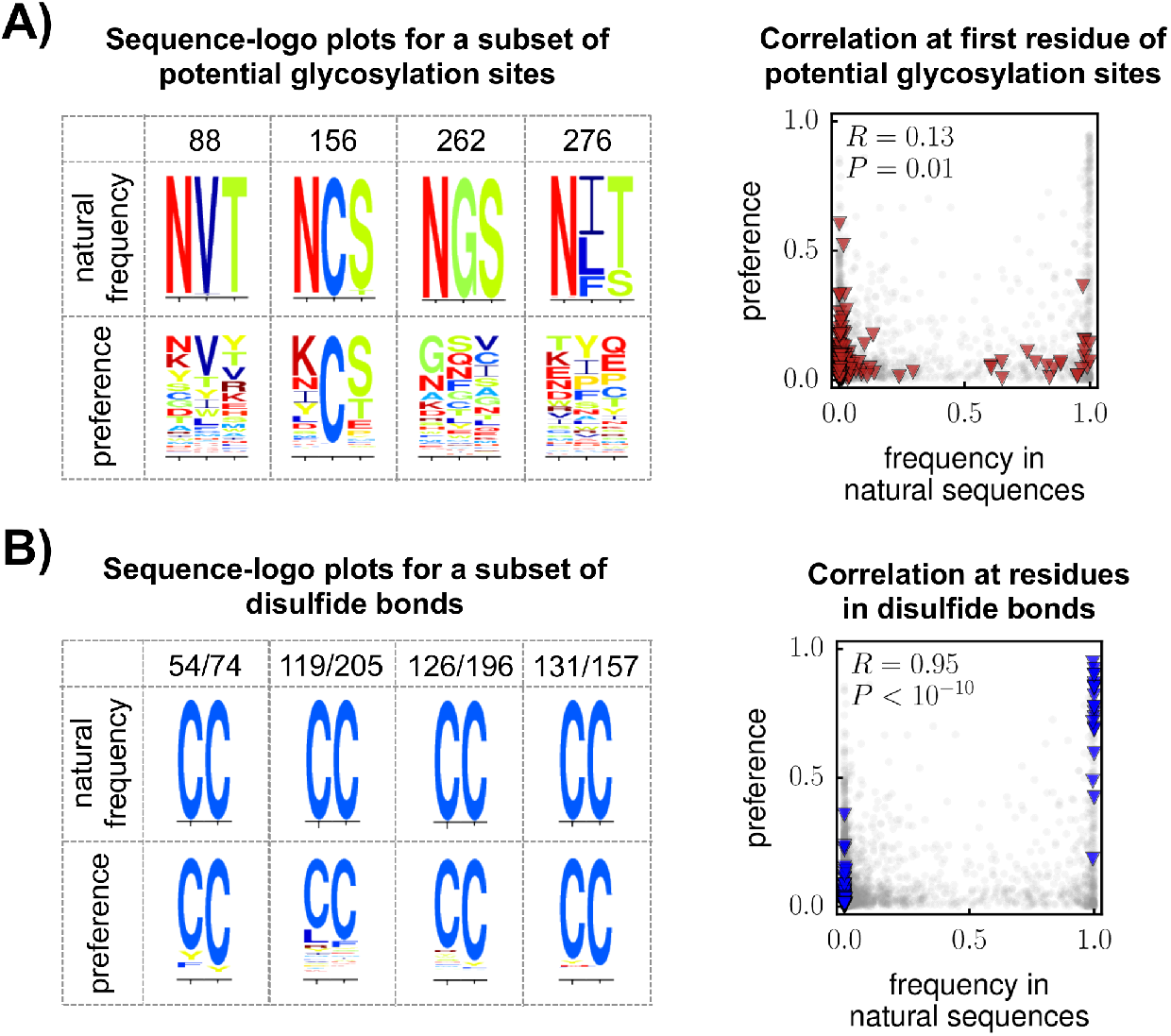
The correlation between the experimentally measured preferences and amino-acid frequencies in natural sequences is low at glycosylation sites, but high at disulfide-bonded cysteines. **(A)** The logo plots show the frequencies of amino acids in the group-M alignment or the amino-acid preferences from our experiments at a subset of potential N-linked glycosylation sites (see S7 Fig for all 30 sites). The glycosylation sites are conserved in nature, but tolerant of mutations in our experiment. The scatter plot shows that there is a poor correlation between the preferences and natural amino-acid frequencies at all 22 alignable glycosylation sites: red triangles represent the first position in each glycosylation site, whereas gray circles represent all other sites. **(B)** There is much better concordance between the preferences and natural amino-acid frequencies for Env’s disulfide-bonded cysteines. The logo plots show each pair of cysteines for a subset of disulfides (see S7 Fig for all 10 disulfides). The scatter plot shows that there is a strong correlation between the preferences and natural amino-acid frequencies at all disulfide-bonded cysteines.

### Env has a high inherent mutational tolerance in its variable loops, and a low mutational tolerance in broadly neutralizing antibody epitopes

Different sites in Env evolve at different rates in natural HIV sequences. These differences arise from two factors. First, some sites are inherently better at tolerating mutations without disrupting Env’s essential functions. Second, some sites are under immune selection that favors rapid change. However, since Env is under selection both to maintain its function and escape immunity, it is difficult to deconvolve these factors.

Our experiments estimate each site’s inherent tolerance for mutations under selection purely for Env’s function in cell culture, without the confounding effects of immune selection (for the remainder of this section, we define a site’s mutational tolerance as the Shannon entropy of its amino-acid preferences shown in Fig 5). We can therefore assess whether regions of Env that evolve rapidly or slowly in nature also have unusually high or low inherent tolerance to mutations. We focused on two regions of Env: the portions of the protein classified as “variable loops” due to extensive variation in nature [77,78], and the epitopes of antibodies that broadly neutralize many HIV strains. We hypothesized that the variable loops would have a high inherent mutational tolerance, whereas broadly neutralizing antibody epitopes would have a low mutational tolerance.

In testing this hypothesis, it is important to control for other properties known to affect mutational tolerance. This can be done by using multiple linear regression to simultaneously analyze how several independent variables affect the dependent variable of mutational tolerance. Relative solvent accessibility (RSA) is the strongest determinant of mutational tolerance in proteins [79], so we included RSA as a variable in the regression. The region of *env* that contains the RRE is under strong nucleotide-level constraint [67–69, Fig 4], so we also included being in the RRE as a binary variable in the regression. We defined the variable loops as indicated in Fig 5, and included being in one of these loops as a binary variable in the regression. Finally, we used crystal structures to delineate broadly neutralizing antibody epitopes. We focused on broadly neutralizing antibodies targeting the CD4 binding site, since most other broadly neutralizing antibodies target either glycans (which are subject to pressures that are not well-modeled in our experiments; Fig 8A) or a membrane-proximal region of gp41 that is not fully resolved in crystal structures of trimeric Env making it impossible to correct for RSA. Specifically, we analyzed the three antibodies with the greatest breadth from [80]: VRC01 (PDB 3NGB [81]), 12A21 (PDB 4JPW [82]), and 3BNC117 (PDB 4JPV [82]). We defined a site as part of an epitope if it was within a 4*Å* inter-atomic distance of the antibody, and included the number of epitopes in which a site is found as a discrete variable in the regression.

The results of the multiple linear regression are in Table 3. As expected, increased solvent accessibility is strongly associated with increased mutational tolerance, whereas presence in the RRE is strongly associated with decreased mutational tolerance. After correcting for these effects, sites in broadly neutralizing epitopes have significantly reduced mutational tolerance. Sites in the variable loops have higher mutational tolerance, although this effect is not statistically significant after controlling for solvent accessibility (mutational tolerance is significantly elevated in the variable loops if solvent accessibility is *not* corrected for; S2 Table). Overall, this analysis provides statistical confirmation of something that is widely assumed in the study of HIV: broadly neutralizing antibodies are unique because they target regions of Env that are inherently intolerant of mutations.

**Table 3.**
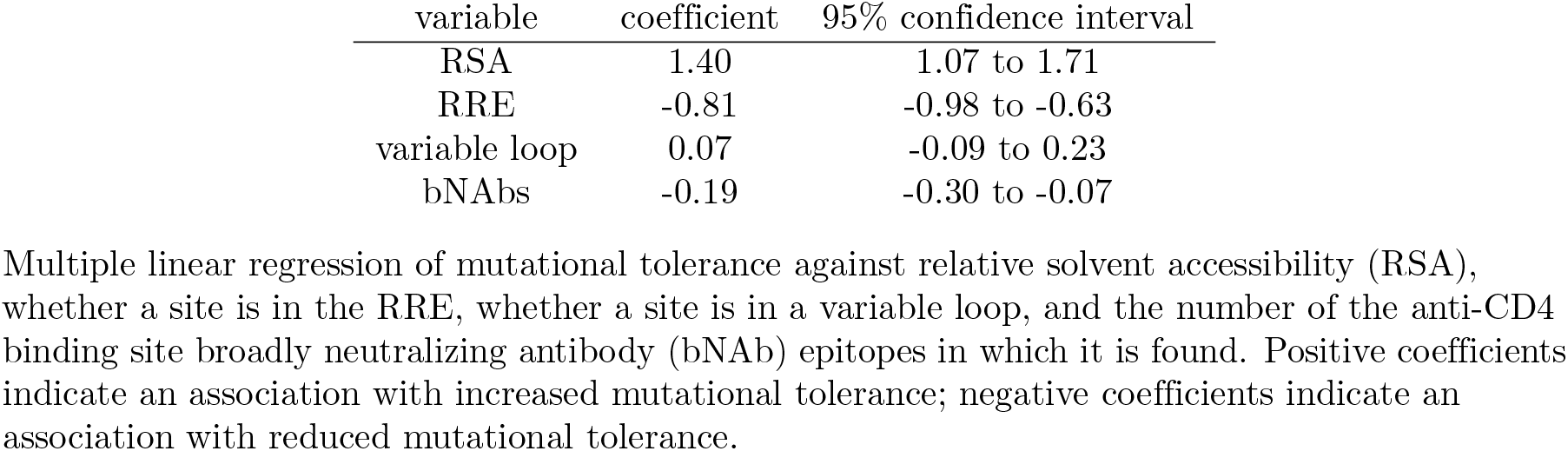
Broadly neutralizing antibody epitopes have significantly lower mutational tolerance than other sites in Env

## Discussion

We have used deep mutational scanning to experimentally estimate the effects of all amino-acid mutations to most of HIV Env. Our experiments select for Env variants that enable HIV to undergo multi-cycle replication in a T-cell line. The broad trends in our data are consistent with what is expected from general considerations of how gene sequence maps to protein function: stop codons are efficiently purged by selection, many but not all nonsynonymous mutations are selected against, and synonymous mutations are less affected by selection except at regions where the nucleotide sequence itself is known to be biologically important. We also find a few sites where nonsynonymous mutations are strongly favored by selection in our experiments, probably because they adapt the virus to cell culture by affecting Env’s conformational dynamics, co-receptor binding, and glycosylation.

We use our experimental data to estimate the preference of each site in Env for each amino acid. We show that these preferences correlate with amino-acid frequencies in natural HIV sequences nearly as well as would be expected if the experimentally measured preferences capture the true selection on Env in nature. The strongest deviations between our measurements and amino-acid frequencies in HIV sequences occur at sites on the surface of the virus that in nature are targeted by pressures (such as antibodies) that are not present in our experiments.

The ability to identify deviations between our measurements and amino-acid frequencies in nature points to a powerful aspect of our approach: it can de-convolve the role of inherent functional constraints and external selection pressures in shaping Env’s evolution. For instance, it is known that some regions of Env regions are conserved in nature, and so can be targeted by broadly neutralizing antibodies. To what extent are these patterns of conservation shaped by Env’s inherent capacity to evolve versus the fact that immune selection targets some parts of the protein more strongly than others? By measuring inherent mutational tolerance at each site under functional selection alone, we show that the epitopes of broadly neutralizing antibodies indeed have a reduced capacity to tolerate mutations irrespective of the action of immune selection.

More generally, our experiments provide high-throughput experimental data that can augment computational efforts to infer features of HIV’s fitness landscape [18–20,22,83]. Such data will aid in efforts to understand viral evolutionary dynamics both within and between patients. Additionally, our experimental approach can be extended to comprehensively examine the effects of mutations on other HIV phenotypes that can be selected in the lab. We anticipate that future experiments could build on the work here to map how mutations affect phenotypes such as cell tropism or sensitivity to antibodies.

## Materials and Methods

### Data and computer code

The computer code to analyze the sequencing data and generate the figures is provided in a series of IPython notebooks in S3 File. Illumina sequencing data are available from the Sequence Read Archive (http://www.ncbi.nlm.nih.gov/sra) under the accession numbers in S10 File.

### Sequence numbering

We use the HXB2 numbering system [44] unless otherwise noted. The “variable loop” definitions were taken from http://www.hiv.lanl.gov/, not including the flanking disulfide-bonded cysteines as part of the loops.

### Codon mutant libraries

We created the codon mutant libraries in the context of the pro-viral genomic plasmid pLAI, which encodes the LAI strain of HIV [40]. This plasmid was obtained from the lab of Michael Emerman. The plasmid sequence is in S4 File.

We created codon mutant libraries of env using the PCR mutagenesis technique described in [31] (see also [32, 37]) except that we performed two total rounds of mutagenesis rather than the three rounds in [31]. The codon tiling mutagenic primers are in S5 File. The end primers were: 5’-ttggaatttctggcccagaccgtctcatgagagtgaaggagaaatatcagcacttg-3’ and 5’-catctgctgctggctcagc-3’. We created three replicate libraries by performing all the steps independently for each replicate starting with independent plasmid preps.

We cloned the PCR mutagenized *env* amplicons into the LAI plasmid with high efficiency to create plasmid mutant libraries. To seamlessly clone the PCR products into the proviral plasmid, we created a recipient version of the plasmid that had *env* replaced by GFP flanked by restriction sites for BsmBI, which cleaves outside its recognition sequence. We named this recipient plasmid pLAI-*δenv*-BsmBI; its sequence is in S6 File. We digested both this recipient plasmid and the gel-purified PCR amplicons with BsmBI (there are BsmBI sites at either end of the PCR amplicon), gel purified the digested PCR products, and ligated them into the plasmid using a T4 DNA ligase. We column purified the ligation products, electroporated them into competent cells (Invitrogen,12033-015), and plated the transformed cells on LB plates supplemented with 100 *μ*g/mL ampicillin. For each of the three replicate libraries, we performed enough transformations to yield >1.4 million unique colonies as estimated by plating dilutions of each transformation on separate plates. Control ligations lacking an insert yielded at least 10-fold fewer colonies. The transformed cells were scraped from the plates, grown in liquid LB-ampicillin at 37°C for ~4 hours, and mini-prepped to obtain the plasmid mutant libraries. For the wildtype controls, we prepped three independent cultures of the wildtype LAI proviral plasmid.

### Generation and passaging of viruses

We generated the mutant virus libraries by transfecting the mutant plasmid libraries into 293T cells. For each replicate, we transfected two 12-well tissue-culture plates to increase the diversity of the generated viruses. Specifically, we plated 293T cells at 2.4×10^5^ cells/well in D10 media (DMEM supplemented with 10% FBS, 1% 200 mM L-glutamine, and 1% of a solution of 10,000 units/mL penicillin and 10,000 *μ*g/mL streptomycin). The next day, we transfected each well with 1 *μ*g plasmid using BioT (Bioland Scientific LLC, B01-01). For the three wildtype controls we used the same process but with only a single 12-well plate per replicate. At one day post-transfection, we aspirated the old media, replacing it with fresh D10. At ~60 hours post-transfection, we filtered the transfection supernatants through 0.4 *μ*m filters. To remove residual plasmid DNA from the transfection, we then treated the filtrate with DNase-I (Roche, 4716728001) at a final concentration of 100 U/mL in the presence of 10 mM magnesium chloride (Sigma, M8266) at 37° C for 20-30 minutes. We froze aliquots of the DNase-treated supernatant at −80°C. Aliquots were thawed and titered by TZM-bl and TCID-50 assays as described below.

We passaged the transfection supernatants in SupT1 cells obtained from the NIH AIDS Reagent Program [84]. SupT1 cells were maintained in a media identical to the D10 described above except that the DMEM was replaced with RPMI-1640 (GE Healthcare Life Sciences, SH30255.01). Before infecting cells, for replicates 1, 2, and 3 (but not replicate 3b), we first filtered thawed transfection supernatants through a 0.2 *μ*m filter in an effort to remove any large viral aggregates. We then infected 10^8^ SupT1 cells with 5 × 10^5^ TZM-bl units of the mutant library transfection supernatant in a final volume of 100 mL SupT1 culture medium in a vented tissue-culture flask (Fisher Scientific, 14-826-80). In parallel, we passaged 10^5^ TZM-bl units of transfection supernatant for each wildtype control in 20 million SupT1 cells in a final volume of 20 mL. At one day post-infection, we pelleted cells at 300× g for 4 minutes and resuspended in fresh media to the same volume as before. At two days post-infection, we added fresh media equal to the volume already in the flask to dilute the cells and provide fresh media. We harvested virus at three days post-infection (for replicates 1, 2, and 3) or four days post-infection (for replicate 3b) by pelleting cell debri at 300×g for 4 minutes and then collecting the viral supernatant for storage at −80°C. To remove residual culture media and plasmid DNA from the cell pellets, we washed pellets two times in PBS. The washed cells were resuspended in PBS to a final concentration of 10^7^ cells/mL, and aliquots were frozen at −80°C for DNA purification.

We conducted a second passage by infecting new cells with the passage-1 viral supernatants. The second passage differed from the first passage in the following ways: Before infecting cells, we filtered passage-1 supernatant of replicate 3b-2 through a 0.2 *μ*m filter but did not filter any of the other replicates. We also had to modify the passaging conditions for some replicates due to low titers of the passage-1 supernatants. For viruses in which the passage-1 supernatant was at too low a concentration to infect at an MOI of 0.005 in the volumes indicated above, we added additional passage-1 supernatant, and then reduced the volume to that indicated above during the day-one media change.

### Virus titering by TCID_50_ and TZM-bl assays

We measured viral titers using TZM-bl reporter cells [85]. Specifically, we added 2× 10^4^ cells in 0.5 mL D10 to each well of a 12-well plate. We made dilutions of viral inoculum and infected cells with 100 uL of each dilution. At 2 days post-infection, we fixed cells in a solution of 1% formaldehyde and 0.2% glutaraldehyde in PBS for 5 minutes at room temperature, washed with PBS to remove the fixing solution, and stained for beta-galactosidase activity with a solution of 4 mM potassium ferrocyanide, 4 mM potassium ferricyanide, and 0.4 mg/mL X-gal in PBS at 37° C for 50 minutes. After washing cells with PBS to remove the staining solution, we used a microscope to count the number of blue cells per well, computing the viral titer as the number of blue cells per mL of viral inoculum.

We were concerned that the infectious titer in SupT1 cells might differ from the TZM-bl titers. We therefore also performed TCID_50_ assay to directly measure infectious titers in SupT1 cells. To do this, we made dilutions of viral transfection supernatant in a 96-well tissue-culture plate and added SupT1 cells at a final concentration of 2.5×10^5^ cells/mL in a final volume of 180 *μ*L/well. At 4 and 8 days post-infection, we passaged supernatant 1:10 into fresh media to prevent cells from becoming over confluent. At 12 days post-infection, we measured the titer of culture supernatants using the TZM-bl assay to determine which SupT1 infections had led to the production of virus. Based on binary scoring from these TZM-bl assays, we calculated titers using the Reed-Muench formula [86] as implemented at https://github.com/jbloomlab/reedmuenchcalculator. At least for the LAI strain used in our experiments, the SupT1 TCID_50_ titers were approximately equal to the TZM-bl titers. Therefore, we used only the less time-consuming TZM-bl assay for all subsequent titering.

### Generation of samples for Illumina sequencing

We purified non-integrated viral DNA from aliquots of frozen cells using a mini-prep kit (Qiagen, 27104) with ~ 10^7^ cells per prep. In some cases, we then concentrated the purified DNA using Agencourt AMPure XP beads (Beckman Coulter, A63880) using a bead-to-sample ratio of 1.0 and eluting with half of the starting sample volume.

We next generated PCR amplicons of *env* to use as templates for Illumina sequencing. We created these amplicons from plasmid or mini-prepped non-integrated viral DNA by PCR using the primers 5’-agcgacgaagacctcctcaag-3’ and 5’-acagcactattctttagttcctgactcc-3’. PCRs were performed in 20 *μ*l or 50 *μ*l volumes using KOD Hot Start Master Mix (71842, EMD Millipore) with 0.3 *μ*M of each primer and 3 ng/*μ*l of mini-prepped DNA or 0.3 ng/*μ*l of plasmid as template. The PCR program was:

1. 95 ° C, 2 minutes
2. 95 °C, 20 seconds
3. 70 ° C, 1 second
4. 64.3 °C, 10 seconds (cooling to this temperature at 0.5 °C/second)
5. 70 ° C, 1 minute 48 seconds
6. Go to 2, 27 times
7. hold at 4 ° C

For replicate 3b, there were a few modifications: the annealing temperature was 64.9 ° C, the extension time was 54 seconds, and we performed only 25 cycles. To quantify the number of unique template molecules amplified in each PCR, we performed standard curves using known amounts of template *env* in pro-viral plasmid, and ran the the bands on an agarose gel alongside our amplicons for visual quantification. We performed a sufficient number of PCR reactions to ensure that amplicons from plasmid were coming from > 10^6^ unique template molecules, and amplicons from viral DNA were coming from ~ 2 × 10^5^ template molecules. All PCR products were purified with Agencourt beads (using a sample-to-bead ratio of 1.0) and quantified by Quant-iT PicoGreen dsDNA Assay Kit (Life Technologies, P7589).

We deep sequenced these amplicons using the strategy for barcoded-subamplicon sequencing in [37], dividing env into six subamplicons (this is a variation of the strategy originally described in [87–89]). The sequences of the primers used in the two rounds of PCR are in S9 File. Our first-round PCR conditions slightly differed from [37]: our 25 *μ*L PCRs contained 12.5 *μ*L KOD Hot Start Master Mix, 0.3 *μ*M of each primer, and 5 ng of purified amplicon. For replicates 1, 2, and 3, the first-round PCR program was:

1. 95 ° C, 2 minutes
2. 95 °C, 20 seconds
3. 70 ° C, 1 seconds
4. 60 °C, 10 seconds (cooling to this temperature at 0.5 °C/second)
5. 70 °C, 10 seconds
6. Go to 2, 10 times
7. 95 °C, 1 min
8. hold 4 °C

For replicate 3b, we used the same program, but with 9 PCR cycles instead of 11. Prior to the second round PCR, we bottlenecked each subamplicon by diluting it to a concentration that should have yielded between 3 and 5 × 10^5^ unique single-stranded molecules per subamplicon per sample. We purified the second-round PCR products using Agencourt beads, quantified with PicoGreen, pooled in equimolar amounts, and purified by agarose gel electrophoresis, excising DNA corresponding to the expected ~500 base pairs in length. We sequenced the purified DNA using multiple runs of an Illumina MiSeq with 2×275 bp paired-end reads.

### Analysis of deep-sequencing data

We used dms_tools (http://jbloomlab.github.io/dms_tools/), version 1.1.dev13, to filter and align the deep-sequencing reads, count the number of times each codon mutation was observed both before and after selection, and infer Env’s site-specific amino-acid preferences using the algorithm described in [70]. The code that performs this analysis is in S3 File. Figures summarizing the results of the deep sequencing are also in this supplementary file.

### Alignment of group-M *env* sequences

We downloaded the 2014 filtered web alignment of env from http://www.hiv.lanl.gov/, including all subtypes for HIV-1/SIVcpz. We then curated this alignment in the following ways. First, we removed sequences differed in length from HXB2 (including gap characters) or contained a premature stop codon, ambiguous residue, or frame-shift mutation. Next, we removed columns in the alignment for which we lacked deep mutational scanning data, columns that had >5% gap characters, or columns in variable loops that appeared poorly aligned by eye. Finally, we randomly selected 30 sequences per subtype for group-M subtypes A, B, C, D, F, and G, for a total of 180 sequences. The resulting alignment is in S7 File. The phylogenetic tree in Fig 7 was inferred using RAxML [90] with the GTRCAT substitution model.

### Computing relative solvent accessibilities

We computed absolute solvent accessibilities based on the PDB structure 4TVP (including all three Env monomers after removing antibody chains) using DSSP [91,92]. We normalized absolute solvent accessibilities to relative ones using the maximum accessibilities provided in the first table of [93]. The relative solvent accessibilities are listed in S8 File.

## Acknowledgments

Thanks to Michael Emerman for providing the LAI plasmid and numerous helpful suggestions, to Julie Overbaugh for numerous helpful suggestions, and to Lily Wu for HIV training.

## Supporting Information

**S1 Table.** Sites of mutations recurrently selected in cell culture.

| sites | error-corrected<br>mutation frequency<br>(P2:DNA) |  |  | wildtype | hydropathy |  |  | RSA | entropy of preferences |
| --- | --- | --- | --- | --- | --- | --- | --- | --- | --- |
|  | 1 | 2 | 3 |  | wildtype | preferences | difference |  |  |
| 48 | 3.4 | 2.1 | 3.4 | A | 1.8 | -0.4 | 2.2 | 0.2 | 2.9 |
| 62 | 1.6 | 4.2 | 8.2 | D | -3.5 | -1.6 | -1.9 | 0.6 | 3.6 |
| 64 | 14.4 | 6.8 | 10.3 | E | -3.5 | -1.6 | -1.9 | 0.6 | 2.8 |
| 65 | 1.3 | 3.3 | 3.1 | V | 4.2 | -2.1 | 6.3 | 0.6 | 3.3 |
| 66 | 6.1 | 3.0 | 13.7 | H | -3.2 | -0.2 | -3.0 | 0.6 | 3.5 |
| 81 | 4.1 | 4.5 | 4.1 | P | -1.6 | -1.9 | 0.3 | 0.6 | 2.9 |
| 105 | 2.2 | 3.1 | 7.0 | H | -3.2 | 0.7 | -3.9 | 0.0 | 3.0 |
| 162 | 11.2 | 4.9 | 1.2 | S | -0.8 | -2.6 | 1.8 | 0.2 | 2.7 |
| 188 | 5.8 | 2.9 | 4.8 | T | -0.7 | -2.1 | 1.4 | 0.5 | 2.9 |
| 203 | -8.6 | 6.7 | 5.4 | Q | -3.5 | 0.0 | -3.5 | 0.0 | 2.5 |
| 207 | 15.6 | 19.2 | 21.1 | K | -3.9 | 2.7 | -6.6 | 0.5 | 2.7 |
| 212 | 5.2 | -27.2 | 9.8 | P | -1.6 | -0.3 | -1.3 | 0.2 | 3.4 |
| 377 | 10.5 | 1.1 | 3.9 | N | -3.5 | -2.7 | -0.8 | 0.2 | 2.2 |
| 420 | 3.0 | 3.8 | 4.8 | I | 4.5 | -1.7 | 6.2 | 0.0 | 2.8 |
| 433 | 7.2 | 7.2 | 8.4 | A | 1.8 | 2.2 | -0.4 | 0.0 | 2.2 |
| 436 | 3.1 | 2.8 | 5.2 | A | 1.8 | 1.1 | 0.7 | 0.0 | 2.4 |
| 443 | -5.4 | 4.2 | 3.2 | I | 4.5 | -0.9 | 5.4 | 0.1 | 3.4 |
| 557 | 4.7 | 11.4 | 5.6 | R | -4.5 | 0.6 | -5.1 | nd | 3.5 |
| 558 | 2.9 | 6.0 | 3.8 | A | 1.8 | -0.7 | 2.5 | nd | 2.2 |
| 560 | 5.9 | 5.2 | 6.0 | E | -3.5 | 0.7 | -4.2 | nd | 3.2 |
| 564 | 42.5 | 6.3 | -4.0 | H | -3.2 | -0.8 | -2.4 | nd | 4.1 |
| 588 | 9.4 | 11.4 | 11.2 | K | -3.9 | 1.5 | -5.4 | 0.2 | 3.0 |
| 591 | 6.2 | 4.2 | 5.0 | Q | -3.5 | 2.2 | -5.7 | 0.0 | 2.3 |
| 653 | 1.6 | 3.4 | 3.2 | Q | -3.5 | 1.7 | -5.2 | 0.5 | 3.1 |
| 655 | 10.8 | 8.2 | 4.7 | K | -3.9 | 1.5 | -5.4 | 0.1 | 3.3 |

**S2 Table.** Mutational tolerance is higher for variable loops when relative solvent accessibility is not taken into account.

| variable | coefficient | 95% confidence interval |
| --- | --- | --- |
| RRE | -0.70 | -0.86 to -0.54 |
| variable loop | 0.16 | 0.01 to 0.31 |
| bNAbs | -0.11 | -0.24 to 0.01 |

Same as Table 3, but not taking RSA into account.

**S1 Fig.**
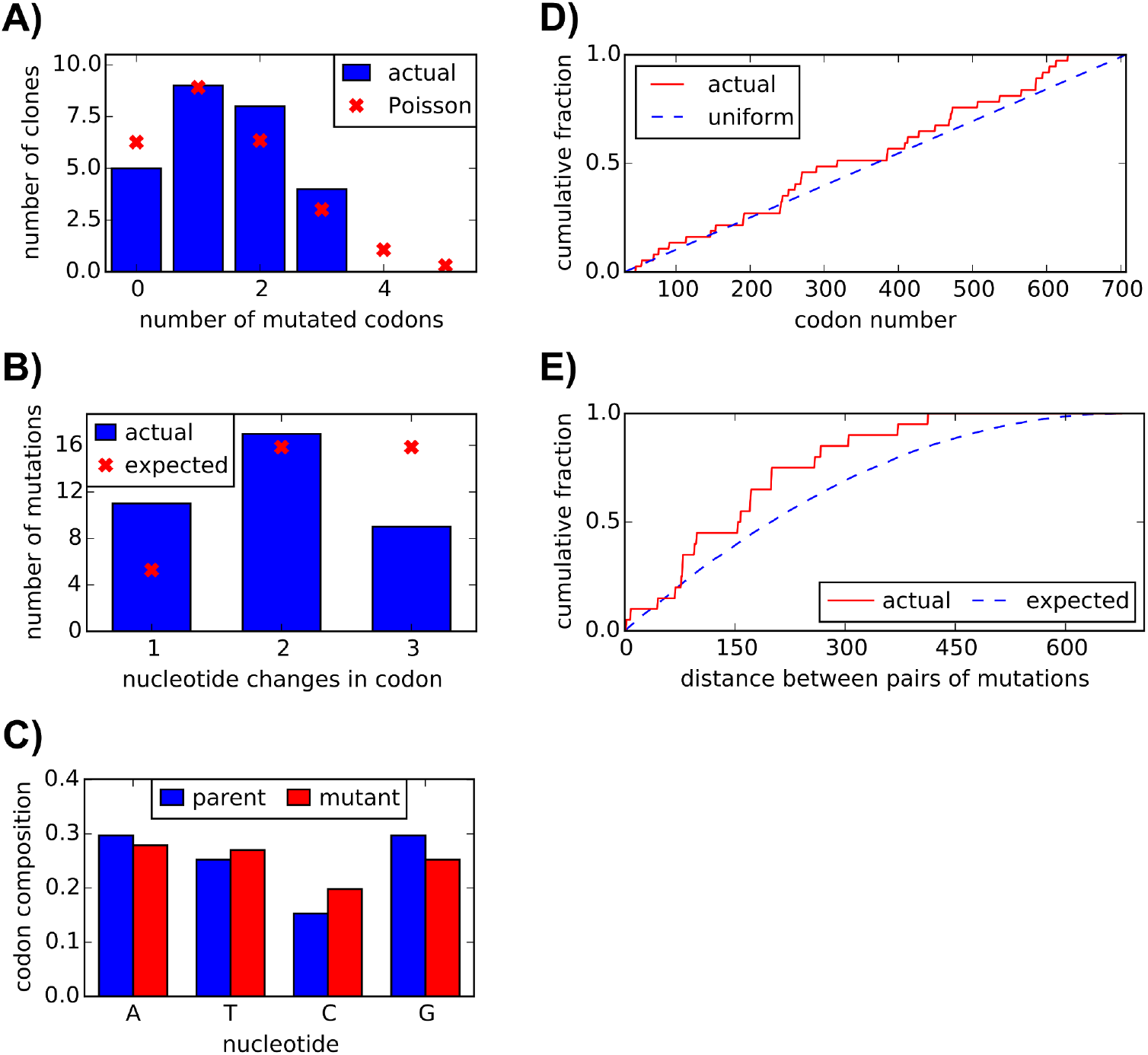
Sanger sequencing of mutant plasmids shows a roughly uniform distribution of codon mutations, with an average of 1.4 mutations per gene. We Sanger sequenced 26 clones sampled roughly evenly from the three replicate mutant plasmid libraries prior to any functional selection. **(A)** We observed an average of 1.4 mutant codons per clone. The number of mutant codons per clone closely followed a Poisson distribution. **(B)** Mutant codons had a mix of single-, double-, and triple-nucleotide changes. **(C)** The nucleotide frequencies were fairly uniform in the mutant codons. **(D)** Mutations were distributed roughly evenly along the portion of env that we mutagenized (codons 31-707). **(E)** For clones with multiple mutations, we computed pairwise distances between mutations in primary sequence and plotted the cumulative distribution of these distances (red line). For comparison, we simulated the expected distribution of pairwise distances if mutations occurred entirely independently (blue line). The difference between the actual and expected distributions suggests our mutagenesis had a slight bias to introduce mutations closer together than expected by chance.

**S2 Fig.**
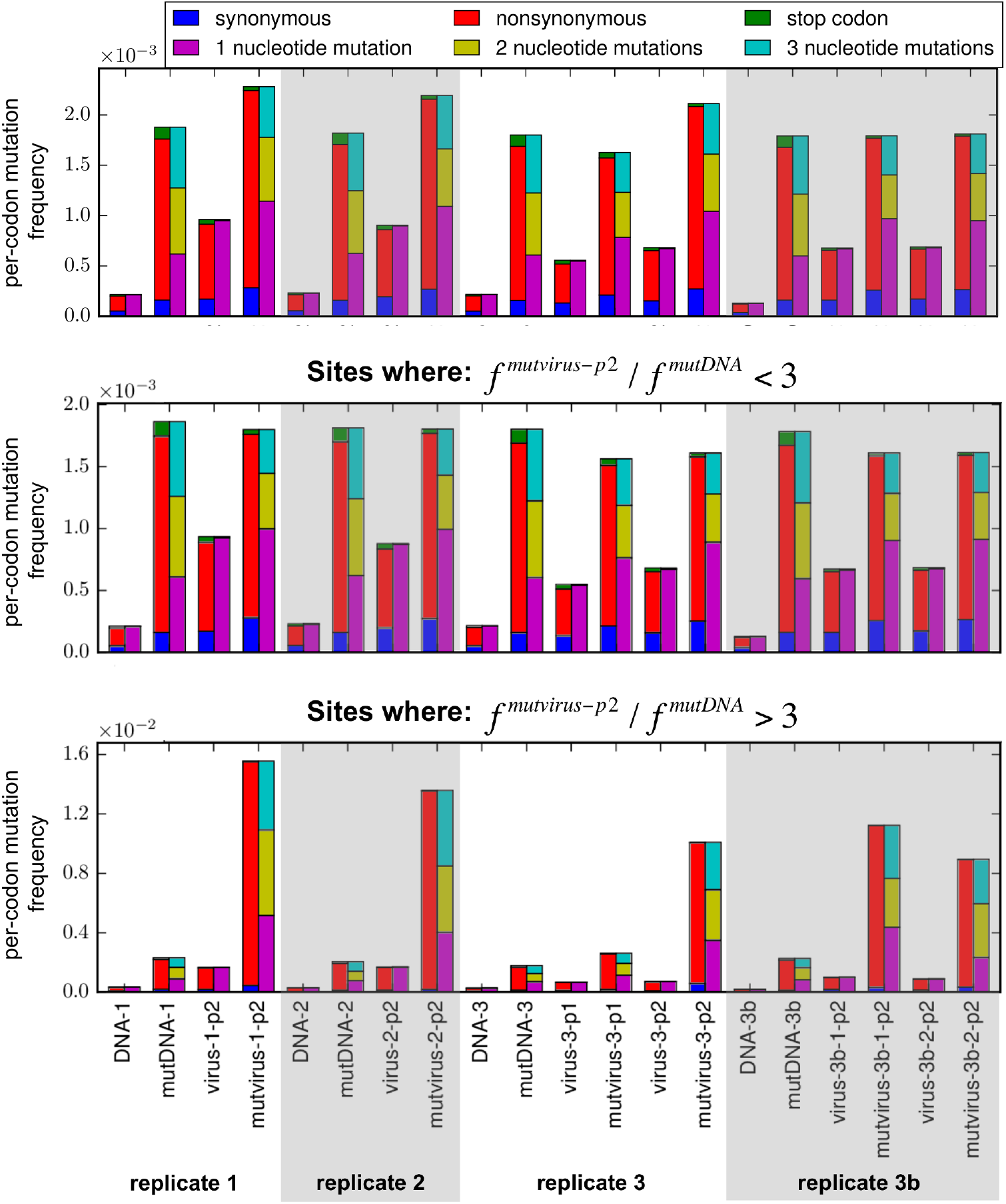
Codon mutation frequencies of mutant libraries and wildtype controls. This figure is similar to Fig 2 except that it shows the uncorrected mutation frequencies in the mutant plasmid and mutant virus libraries, and the mutation frequencies in the wildtype controls that were used to correct the mutation frequencies in Fig 2. Codon mutations are classified both by their effect on the protein (synonymous, nonsynonymous, or stop codon) and by the number of nucleotides they change in the codon (one, two, or three). The top panel shows data for all sites, whereas the middle and lower panels show data for the indicated subsets of sites. This this plot, *f^mutmrus–p2^* and *f^mutDNA^* refer to the nonsynonymous mutation frequency in the twice-passaged mutant viruses and the initial mutant DNA, respectively.

**S3 Fig.**
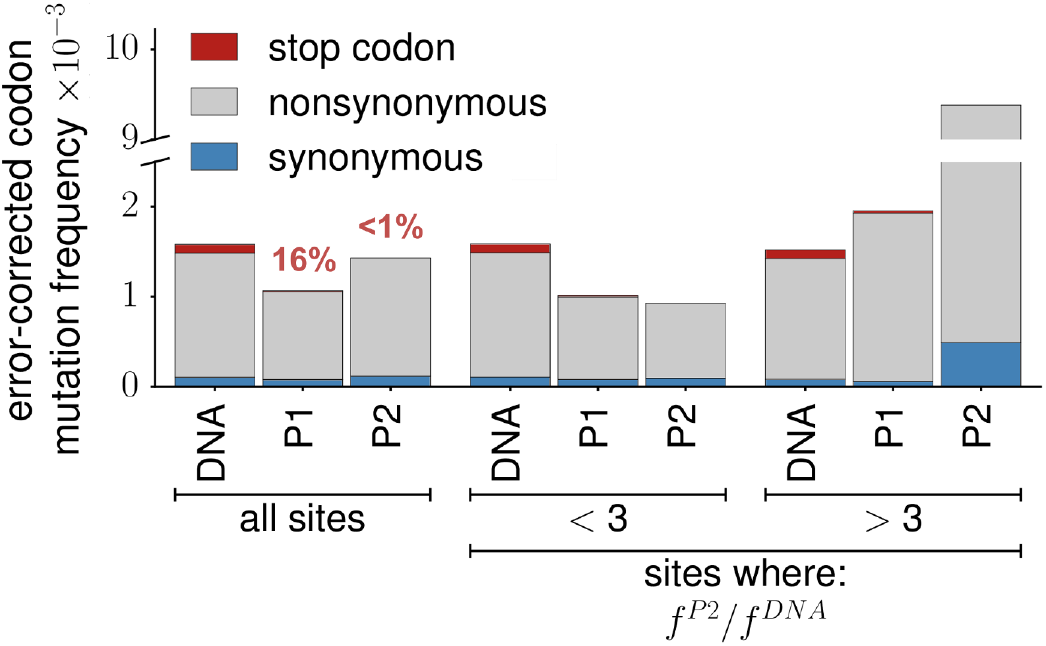
Complete selection against stop codons requires two rounds of viral passage. We deep sequenced the replicate 3 library after both one (P1) and two (P2) rounds of viral passaging. This figure is similar to Fig 2, but shows data for both P1 and P2. Purging of stop-codon mutations shows selection was only complete after two rounds of passaging. Whereas two rounds of passaging purged stop-codon mutations to <1% their frequency in the initial library (DNA), one round of passaging only purged stop-codon mutations to 16% their starting frequency (see the data for “all sites”, where the red numbers above the bars for P1 and P2 indicate the percentage of stop codons after each passage relative to the starting library).

**S4 Fig.**
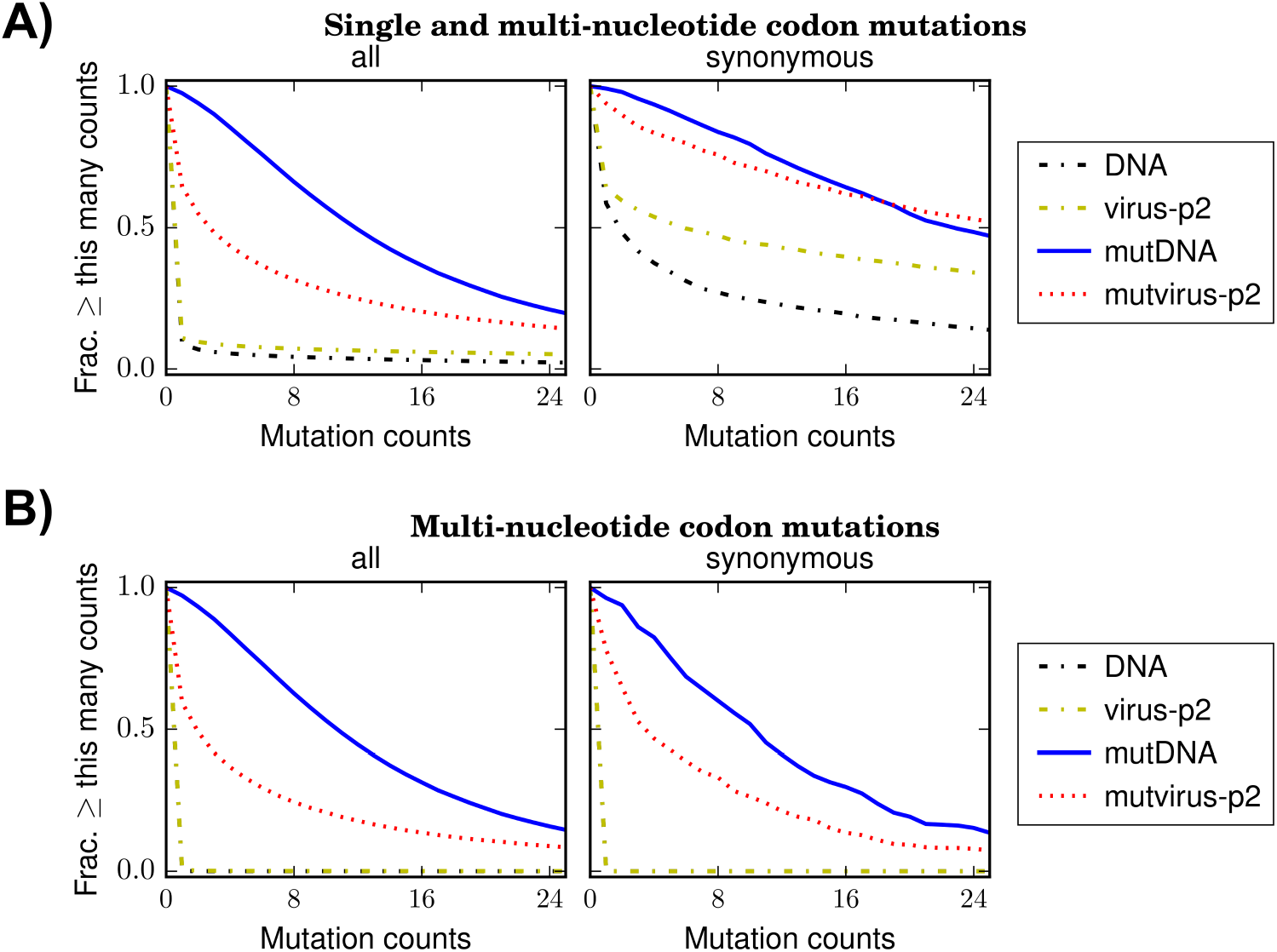
Sampling of codon mutations in all replicates combined. **(A)** Each plot shows the number of all (single and multi-nucleotide) codon mutations observed at least the indicated number of times in the sequencing of all replicates combined. We observed almost all mutations in the starting plasmid libraries (mutDNA), showing rich initial mutational diversity. Many mutations were depleted in the mutant virus libraries after two rounds of passaging (mutvirus-p2), consistent with purifying selection purging deleterious variants or bottlenecking diminishing library diversity. Examination of mutation counts in the wildtype plasmid (DNA) and wildtype virus (virus-p2) controls revealed a considerable fraction of mutations that were present at appreciable numbers due to errors from deep sequencing and PCR or de *novo* mutations from viral replication. This observation underscores the importance of using these wildtype controls to correct for background errors and de novo mutations. **(B)** If we examine only multi-nucleotide codon mutations, then there are negligible background errors in the wildtype controls. Similar data for each replicate individually are in S5 Fig.

**S5 Fig.**
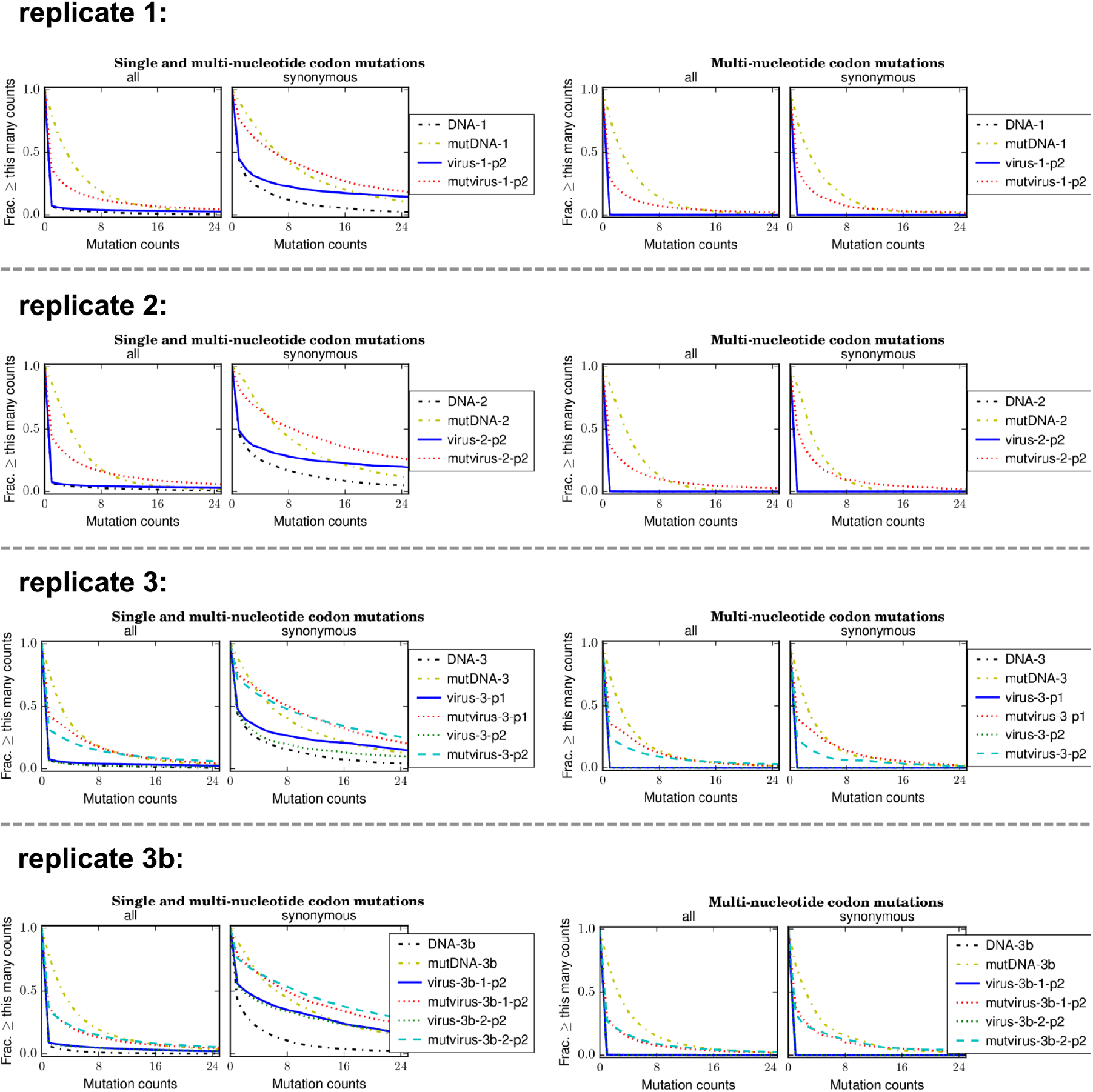
Sampling of codon mutations in individual replicates. These plots are the same as in S4 but show each replicate individually.

**S6 Fig.**
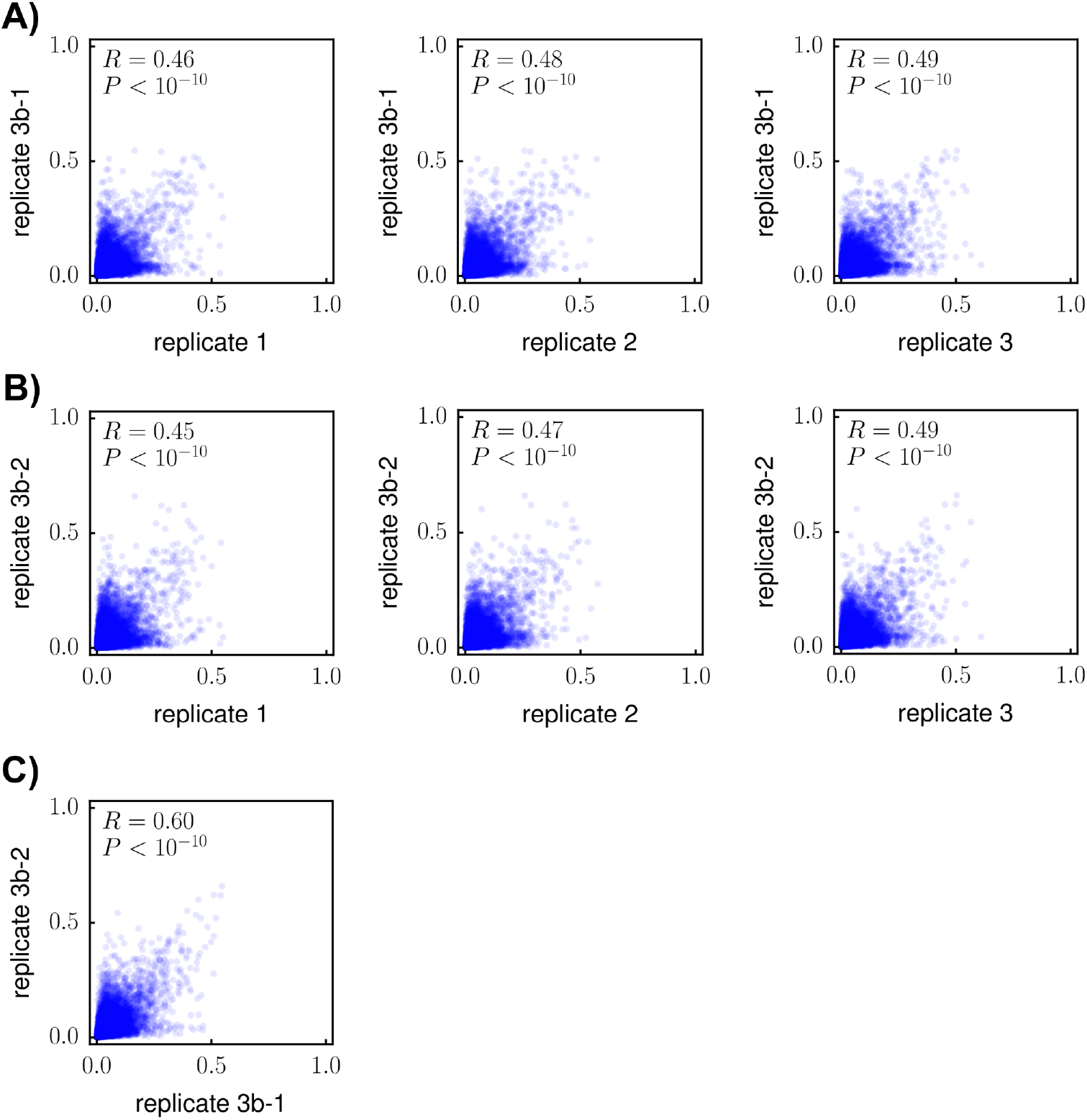
Correlation of site-specific amino-acid preferences between replicates, including 3b-1 and 3b-2. **(A)** The correlation between replicate 3b-1 and replicate 1, 2, or 3. **(B)** The correlation between replicate 3b-2 and replicate 1, 2, or 3. **(C)** The correlation between replicates 3b-1 and 3b-2.

**S7 Fig.**
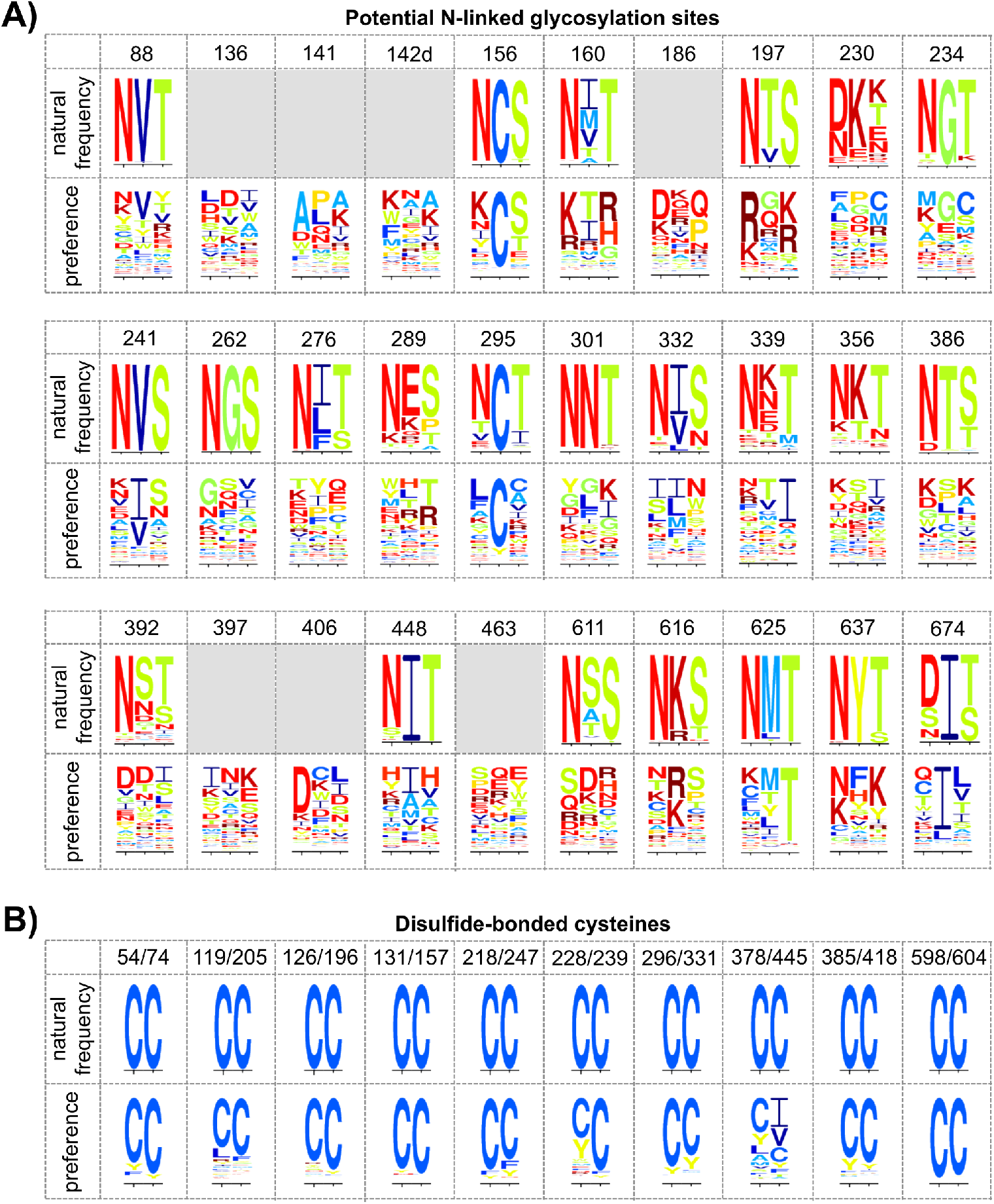
Amino-acid frequencies and preferences for all potential N-linked glycosylation sites and disulfide bonds. This figure is similar to Fig 8, but shows logo plots for all 30 glycosylation sites (defined using the N-GlycoSite tool [100] from the HIV sequence database, http://www.hiv.lanl.gov/) and all 10 disulfide bonds [101] in LAI. **(A)** Most glycosylation sites are highly conserved in natural sequences, but highly tolerant of mutations in our experiments. Logo plots showing amino-acid frequencies in nature are replaced by grey boxes for sites in the alignment of group-M sequences that were masked because the site had >5% deletions relative to HXB2 or because the region looked unalignable by eye (for details, see IPython notebook `CurateLANLMultipleSequenceAlignment.ipynb` within S3 File). **(B)** Disulfide-bonded cysteines are absolutely conserved in nature, and most have a strong preference for cysteine in our experiments. A previous study [43] found that only the C378-C445 disulfide bond tolerated alanine mutations at individual cysteines while supporting robust viral replication in cell culture. In accordance with this previous work, these cysteines are the most mutationally tolerant ones in our experiment.

## All Supplementary Files are included in a single ZIP archive

**S1 File. Average of the amino-acid preferences measured in the replicates**. Sites are numbered using the HXB2 scheme. The same preferences re-scaled by the optimal stringency parameter are in S2 File.

**S2 File. Amino-acid preferences re-scaled by the optimal stringency parameter**. The preferences in S1 File re-scaled by the optimal stringency parameter of *β* = 2.1. These are the data plotted in Fig. 5. However, preferences for stop codons are listed in this file, but not shown Fig 5.

**S3 File**. iPython notebooks that perform the data analysis steps described in this paper.

**S4 File**. A Genbank file with the sequence of the LAI pro-viral plasmid.

**S5 File**. The codon tiling primers used to construct the mutant libraries.

**S6 File**. The recipient pro-viral plasmid, which has *env* replaced by partial GFP and beta globin genes flanked by BsmBI sites.

**S7 File**. The alignment of group M Env sequences.

**S8 File**. The relative solvent accessibilities of all sites in Env present in the crystal structure.

**S9 File**. The PCR primers used in the barcoded-subamplicon sequencing.

**S10 File**. The SRA accession numbers for deep sequencing data. Samples are named as follows: mutDNA-1 denotes the mutant plasmid library for replicate 1; DNA-1 denotes the wildtype plasmid for replicate 1; mutvirus-p2-1 denotes the twice-passaged mutant viral libraries for replicate 1; virus-p2-1 denotes the twice-passaged wildtype virus for replicate 1.

